# A hydrophobic core in the coiled-coil domain is essential for NRC resistosome function

**DOI:** 10.1101/2025.01.21.634219

**Authors:** Hung-Yu Wang, Enoch Lok Him Yuen, Kim-Teng Lee, Foong-Jing Goh, Tolga O. Bozkurt, Chih-Hang Wu

## Abstract

The nucleotide-binding leucine-rich repeat protein (NLR) required for cell death (NRC) family represents a group of helper NLRs that are required by sensor NLRs to execute hypersensitive cell death during pathogen infection. NRCs contain an N-terminal coiled-coil (CC) domain essential for their function, yet our knowledge of how this domain contributes to NRC function remains limited. Here, we identified a novel hydrophobic feature within the CC domain that contributes to NRC-mediated immunity. We screened for conserved hydrophobic residues among NRCs and identified seven required for NRC4-mediated cell death. Structural analysis revealed that four of these residues form a hydrophobic core in the CC domain. This hydrophobic core is important for NRC4 subcellular localization, oligomerization, and phospholipid association, but not for NRC4 focal accumulation at the extrahaustorial membrane during *Phytophthora infestans* infection. Sequence analysis and functional assays revealed this core is highly conserved in NRCs and some singleton NLRs but has degenerated in NRC-dependent sensor NLRs. Our study identifies a novel hydrophobic feature in the CC domain of NRCs and reveals its contribution to NLR-mediated immunity.

## Introduction

Plants, as immobile organisms, have evolved sophisticated immune receptor systems to detect and respond to pathogen attacks (Jones & Dangl, 2006; Ngou *et al*, 2021; Jones *et al*, 2024). Among these defense mechanisms, nucleotide-binding leucine-rich repeat proteins (NLRs) serve as intracellular immune sensors that recognize pathogen effectors and initiate hypersensitive cell death to restrict pathogen proliferation (Jones & Dangl, 2006; Duxbury *et al*, 2021).

NLRs consist of three canonical domains: the N-terminal domain, the NB-ARC domain, and the LRR domain (Duxbury *et al*, 2021). Based on their N-terminal domains, NLRs are classified into three main types: CC-NLRs (CNLs) containing a coiled-coil domain, TIR-NLRs (TNLs) with a Toll/interleukin-1 receptor domain, and CC_R_-NLRs (RNLs) possessing a Resistance to Powdery Mildew 8 (RPW8)-like domain (Duxbury *et al*, 2021). Different N-terminal domains mediate distinct downstream events leading to NLR-triggered immunity. In TNLs, the TIR domain functions as an NADase enzyme, producing phosphoribosyl adenosine monophosphate/diphosphate (pRib-AMP/ADP) or cyclic ADPR (cADPR) isomers that initiate immune signaling (Ma *et al*, 2020; Martin *et al*, 2020; Huang *et al*, 2022; Jia *et al*, 2022; Yu *et al*, 2024). Mutations in the enzymatic residues essential for NADase activity disrupt immune signaling and impair TNL function (Wan *et al*, 2019; Ma *et al*, 2020). Recent studies on CNLs and RNLs have demonstrated that their CC domains play crucial roles in membrane association, with these proteins likely functioning as calcium-permeable channels upon activation to trigger cell death (Wang *et al*, 2019a; Bi *et al*, 2021; Jacob *et al*, 2021). Mutations in positively charged residues involved in membrane association or negatively charged residues critical for ion channel activity disrupt cell death mediated by some CNLs and RNLs (Bi *et al*, 2021; Jacob *et al*, 2021; Saile *et al*, 2021; Wang *et al*, 2023).

While some NLRs like ZAR1 act as functional singletons that both recognize effectors and initiate immune responses, others have undergone functional specialization, evolving into sensor NLRs that detect effectors and helper NLRs that activate immune responses (Wu *et al*, 2017, 2018; Adachi *et al*, 2019b; Contreras *et al*, 2023a; Shepherd *et al*, 2023). The NLR-required for cell death (NRC) family represents a group of helper NLRs that form a complex network with phylogenetically related sensor NLRs, particularly expanded in asterid plants (Wu *et al*, 2017; Goh *et al*, 2024; Sakai *et al*, 2024). In solanaceous plants, NRC2, NRC3, and NRC4 are the three major NRCs that function with various sensor NLRs, forming a complex immune network against a broad range of pathogens (Wu *et al*, 2017). Some sensor NLRs rely on a single NRC for immunity signaling, while others utilize multiple NRCs, highlighting the genetic redundancy within the NRC network. For example, the sensor NLR *Rpi-blb2*, which detects the RXLR effector AVRblb2 delivered by *Phytophthora infestans*, signals through NRC4, whereas another sensor NLR, Rx, which detects the coat protein of potato virus X, signals through NRC2/3/4 in *Nicotiana benthamiana* (Wu *et al*, 2017).

Recent cell biology and cryo-EM studies have revealed that NRCs transition from cytosolic dimer complexes in their resting state to membrane-associated hexameric resistosome complexes upon activation (Madhuprakash *et al*, 2024; Liu *et al*, 2024; Selvaraj *et al*, 2024; Ma *et al*, 2024). In the resting state, NbNRC2 protomers form homodimers with unique dimerization interfaces in the cytosol that likely prevent unwanted cross-activation between different NRC helpers (Selvaraj *et al*, 2024). Additionally, studies on tomato NRC2 (SlNRC2), an ortholog of NbNRC2, revealed that, in addition to homodimers, SlNRC2 assembles into filamentous structures that likely serve to regulate its activity negatively (Ma *et al*, 2024). Upon activation by sensor NLRs, the NbNRC homodimers dissociate and oligomerize into hexameric resistosome complexes on the plasma membrane (Wang *et al*, 2023; Liu *et al*, 2024). Notably, NbNRC4 resistosomes function as ion-permeable channels that enable calcium influx, ultimately inducing hypersensitive cell death (Liu *et al*, 2024). This role of hexameric NRC resistosomes aligns with findings from the pentameric AtZAR1 and TmSr35 resistosomes, which also form ion channels on the plasma membrane to trigger cell death (Wang *et al*, 2019a; Liu *et al*, 2024; Förderer *et al*, 2022).

During *P. infestans* infection, the pathogen forms haustoria that penetrate plant cells to deliver effectors and obtain nutrients (Wang *et al*, 2017; Bozkurt & Kamoun, 2020). These haustoria are surrounded by a specialized, newly-synthesized plant membrane called the extrahaustorial membrane (EHM) (Bozkurt & Kamoun, 2020; Yuen *et al*, 2023). To successfully colonize the host, *P. infestans* secretes a range of RXLR effectors across the EHM to facilitate the infection (Bozkurt *et al*, 2011; Wang *et al*, 2019b; King *et al*, 2024). Among these, the RXLR effector AVRblb2 accumulates at the EHM and targets the secretion of papain-like cysteine protease C14, thereby suppressing immune responses (Bozkurt *et al*, 2011, 201). Interestingly, NRC4 focally accumulates at the EHM during infection, even in the absence of its cognate sensor NLR Rpi-blb2 (Duggan *et al*, 2021). Upon activation, NRC4 forms puncta on both the EHM and the plasma membrane. Recent research has highlighted the importance of interactions between NRC4 and phosphatidylinositol phosphates, such as PI4P, which are also present on the EHM (Rausche *et al*, 2021; Wang *et al*, 2023). However, the extent to which NRC4-lipid interactions influence its localization at the EHM and the plasma membrane remains to be determined.

As a member of the CNL family, NRC4 contains four α helices in its CC domain (Liu *et al*, 2024). The first α helix harbors a conserved MADA motif, which is shared by NRC helper proteins and ZAR1 but not by NRC-dependent sensor NLRs (Adachi *et al*, 2019a). Mutations in the hydrophobic residues of the MADA motif disrupt NRC and ZAR1-mediated cell death, highlighting the importance of hydrophobicity within this region for NRC function (Adachi *et al*, 2019a). However, the contributions of the other α helices in the CC domain to NRC4 function remain unknown.

In this study, we identified a novel hydrophobic core formed by the α2–α4 helices of the NRC4 CC domain which plays a critical role in NRC4-mediated immunity. Mutations in this hydrophobic core impaired NRC4 subcellular localization, resistosome formation, and NRC4-phospholipid association without affecting its focal accumulation at the EHM during *P. infestans* infection. Further investigation revealed that the corresponding residues in this hydrophobic core are highly conserved in NRC helper NLRs and some singleton NLRs like ZAR1 and Rpi-vnt1.1 but have diverged in NRC-dependent sensor NLRs. Moreover, while this hydrophobic feature is functionally important in ZAR1, Rpi-vnt1.1, and NRCs, it appears to have degenerated in sensor NLRs. Our findings reveal a previously uncharacterized hydrophobic element at the center of the CC domain that is essential for NRC4 subcellular localization, resistosome formation, and phospholipid binding, and is conserved across NRC helper NLRs and some singleton NLRs, providing new insights into the structural basis of NLR-mediated immunity in plants.

## Results

### Mutations of conserved hydrophobic residues in the CC domain compromise the cell death activity of NRC4

The N-terminal CC domain in CNLs comprises four alpha helices (α1∼α4) crucial for resistosome function (Wang *et al*, 2019a; Liu *et al*, 2024). In NRC4 and most other NRCs, the α1 helix contains a MADA motif essential for mediating cell death (Adachi *et al*, 2019a). However, how α2-4 helices in the CC domain contribute to the NRC function is not clear. To dissect the function of these alpha helices in NRC4-mediated cell death, we perform a full-length protein sequence alignment of NRC4 with NRC2, NRC3, and AtZAR1. We identified 19 conserved hydrophobic residues across the three alpha helices of NRCs and AtZAR1 (Fig. S1). To determine which residues are essential for NRC4 function, we mutated those hydrophobic residues (L/A/I/V/F) to the hydrophilic residue glutamic acid (E). We then tested their cell death activities by co-expression with Rpi-blb2 and AVRblb2 in the *nrc2/3/4*_KO *N. benthamiana* and quantified the cell death by using auto-fluorescence-based imaging (Fig. 1A). Among the 19 NRC4 variants, we found that single mutations at L34, F44, A72, I114, V118, L121, and L128 fully impaired NRC4-mediated cell death (Fig. 1B). Protein accumulation levels of these NRC4 mutants with compromised cell death showed no significant difference compared to the wild-type (Fig. S2A, B), indicating that the reduced NRC4-mediated cell death was not due to changes in protein stability. In total, we identified mutations in 2 residues in α2, 1 residue in α3, and 4 residues in α4 of the CC domain that compromised NRC4-mediated cell death.

**Figure 1.**
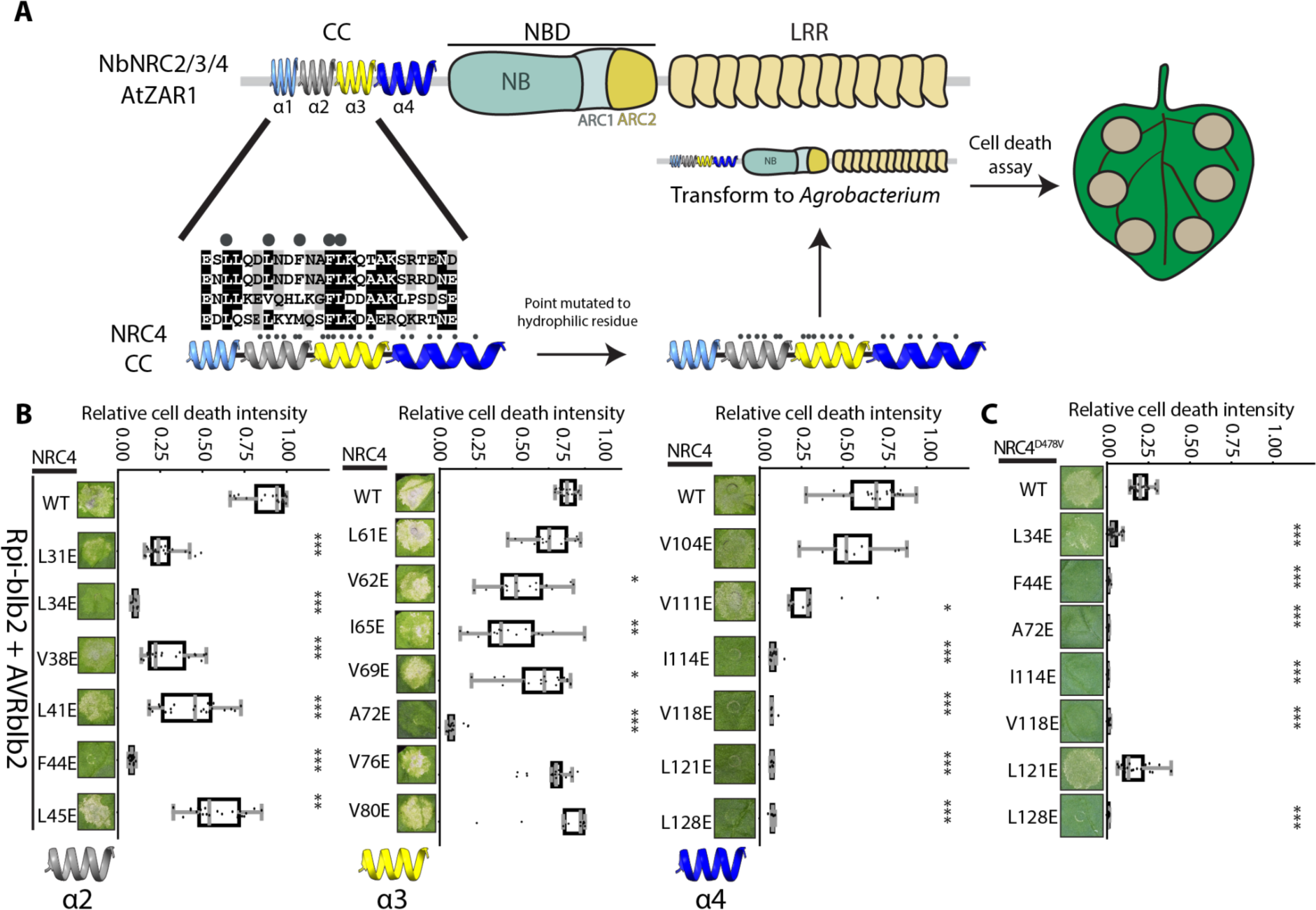
Mutation of conserved hydrophobic residues in the CC domain affects the cell death activity of NRC4. (A) Experimental design for screening key hydrophobic residues in the NRC4 CC domain. Conserved hydrophobic residues shared by NRC2/3/4 and AtZAR1 were selected and mutated to glutamic acid. The generated NRC4 variants were then tested in cell death assays. (B) Cell death assays of NRC4 hydrophobic variants, co-expressed with Rpi-blb2 and AVRblb2 in *nrc2/3/4*_KO *N. benthamiana*. Cell death intensity and phenotypes were recorded at 6 days post-infiltration (dpi). (C) Cell death assays of autoactive NRC4 variants with candidate hydrophobic residue mutations identified in panel B. For all cell death assays, the central line in each box plot represents the median cell death intensity, with box edges indicating the 25th and 75th percentiles. The whiskers extend to the most extreme data points no more than 1.5 x of the interquartile range. Statistical differences between wild-type and variant samples were analyzed using the paired Wilcoxon signed rank test (*=p<0.01, **=p<0.005, ***=p<0.00001).

To test whether these 7 mutations also compromise the cell death induced by an autoactive NRC4 mutation, we introduced them individually into the NRC4^D478V^, which contains a mutated MHD motif leading to sensor-independent cell death. Most of these mutations also completely impair NRC4^D478V^-mediated cell death, with the exception of L121E. While the NRC4^L121E^ mutation failed to induce cell death when co-expressed with Rpi-blb2/AVRblb2, NRC4^L121E/D478V^ induced strong cell death (Fig. 1C). This suggests that residue L121 may play a role in the cooperation between NRC4 and its sensor NLRs. As we decided to focus on intrinsic features of the NRC4 CC domain, we excluded L121 from further analyses. In summary, we identified six key hydrophobic residues in helices α2 to α4 of the CC domain that contribute to NRC4-mediated cell death.

### The identified residues form a hydrophobic Core in the CC domain

Using NRC2 (PDB: 8RFH) as a reference, we employed Alphafold3 (AF3) to model the dimer structure of NRC4, including the CC domain that was not captured in the original NRC2 cryo-EM data (Fig. 2, S3, and S4). We discovered that four of the identified residues (L34/A72/I114/V118) are located in proximity to each other, likely forming a hydrophobic core within the CC domain (Fig. 2A and S3). In contrast, the two other residues, F44 of α2 and L128 of α4, are positioned near each other but distant from the hydrophobic core (Fig. 2A).

**Figure 2.**
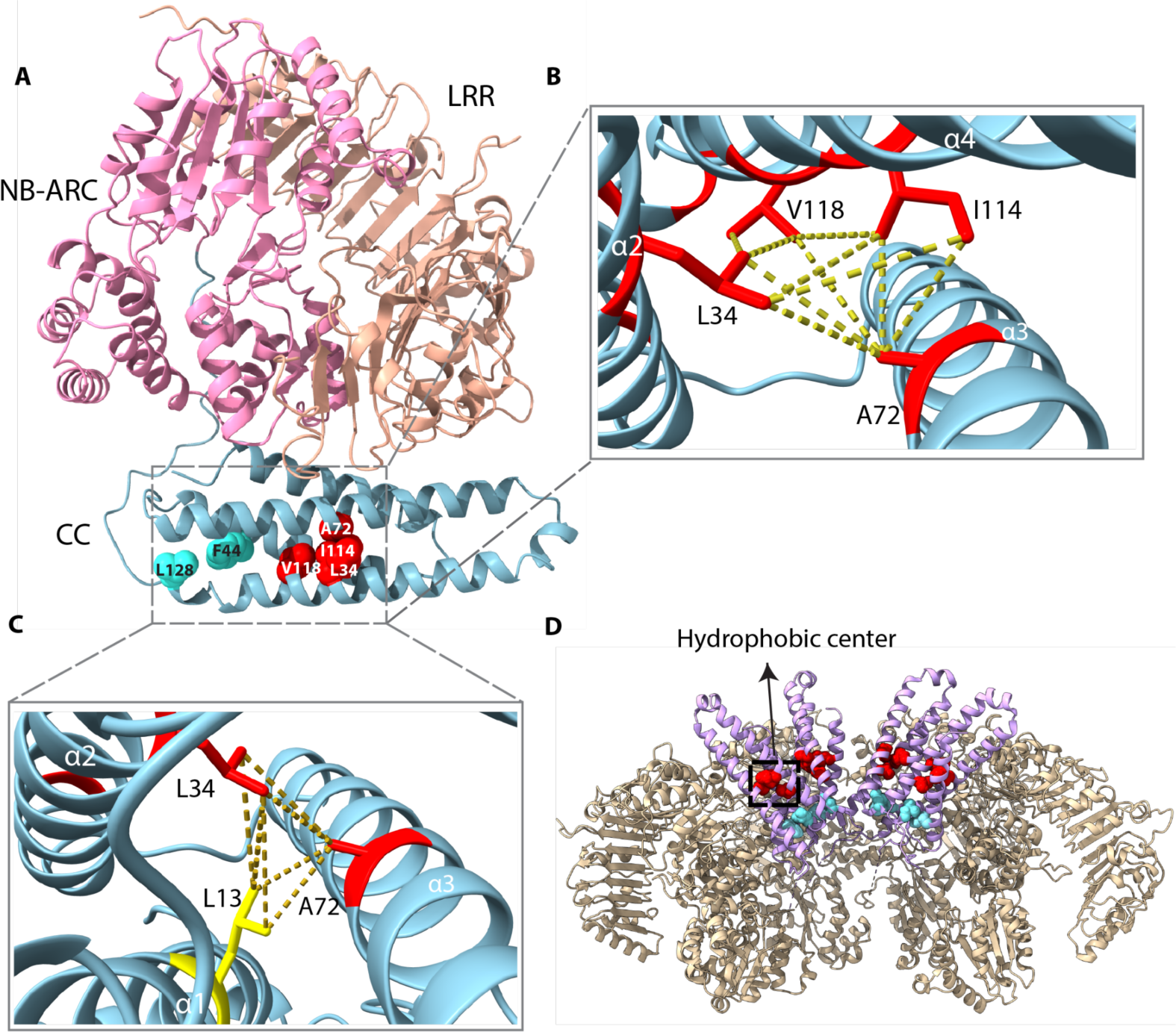
Four of the identified residues form a hydrophobic core at the center of the NRC4 CC domain. (A) The predicted NRC4 structure at the resting state. The coiled-coil (CC) domain is shown in light blue, the NB-ARC domain in pink, and the LRR domain in beige. The four residues that form the hydrophobic core are highlighted in red. The two other residues that are critical for NRC4 function but distant from the hydrophobic core, F44 and L128, are highlighted in cyan. (B) Close-up view of the hydrophobic core formed by L34, A72, I114, and V118 (red). Dashed lines indicate interactions where side-chain distances are less than 6 Å. (C) Spatial relationship between the hydrophobic core residues (red) and L13 (yellow) of the MADA motif in the α1 helix. Dashed lines represent potential interactions where side-chain distances are less than 6 Å. (D) The location of the hydrophobic core within the NRC4 resistosome (PDB: 9CC8). For clarity, only four of the six NRC4 protomers are displayed. The CC domains are shown in purple, with the remaining regions in gold. Hydrophobic core residues are highlighted in red, while F44 and L128 (residues outside the core) are indicated in cyan. All molecular structures (AlphaFold3-predicted models or PDB structures) were visualized using ChimeraX.

**Figure 3.**
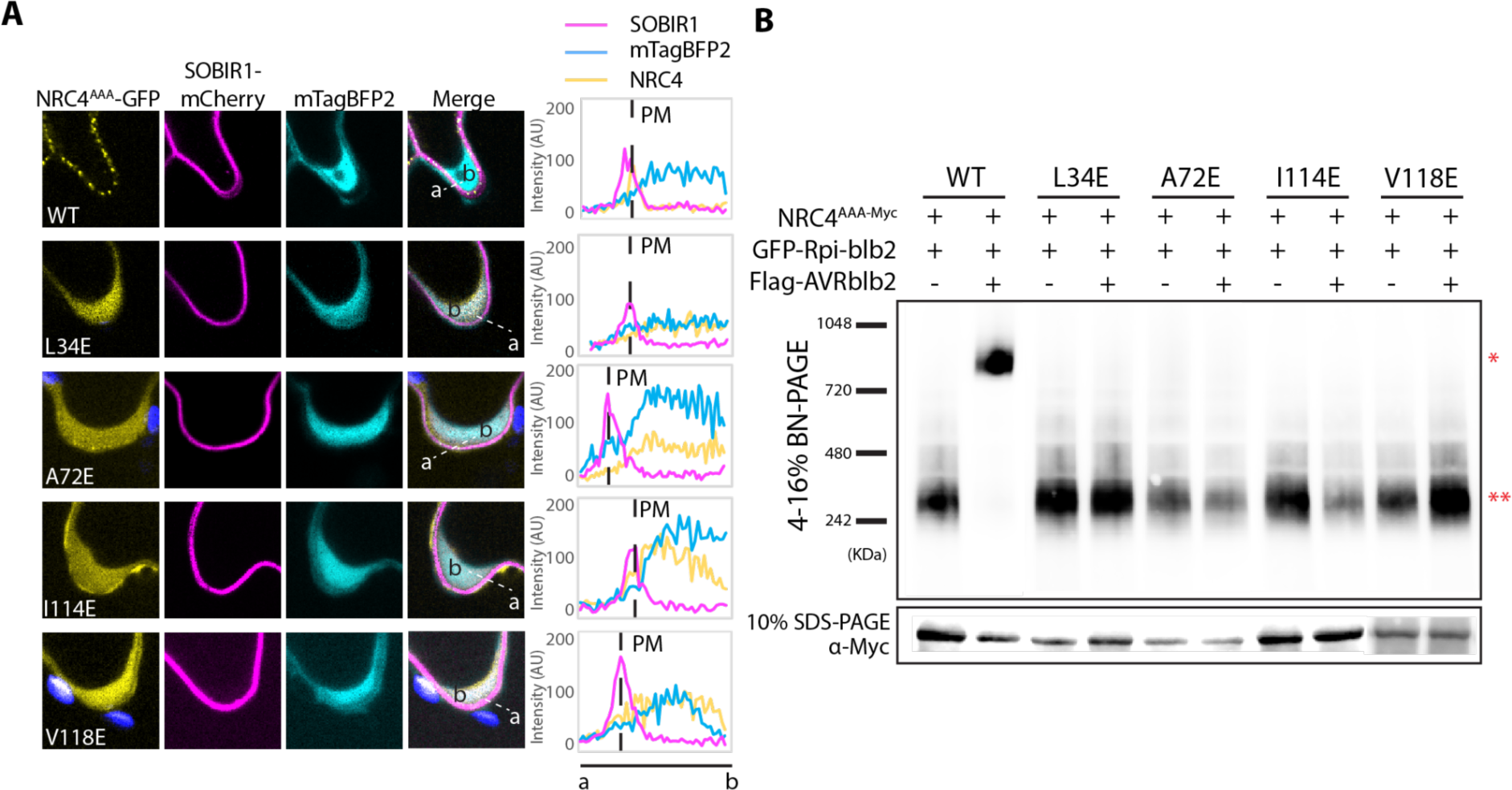
Mutations in the hydrophobic core disrupted NRC4 puncta formation and oligomerization. (A) Confocal images showing subcellular localization of transiently-expressed GFP-tagged NRC4^AAA^ and hydrophobic core mutants co-expressed with SOBIR1-mCherry (plasma membrane marker) and mTagBFP2 (cytosol marker) in *nrc2/3/4*_KO *N. benthamiana* leaves that also co-express Rpi-blb2 and AVRblb2. Images were captured at 2 dpi. The lines in the overlay panel indicate the region selected for measuring the fluorescence intensity of each channel. The blue signal corresponds to the chloroplast. The intensity was measured from position “a” to “b,” and values are reported in arbitrary units (AU). (B) Upper panel: BN-PAGE analysis showing the formation of the NRC4^AAA^ and hydrophobic core mutant resistosomes. NRC4 variants and Rpi-blb2 were transiently expressed in *nrc2/3/4*_KO *N. benthamiana* leaves, with and without AVRblb2. Lower panel: Western blot analysis of SDS-PAGE to confirm NRC4 protein expression levels. Total protein samples were collected at 2 dpi for both BN-PAGE and SDS-PAGE. Approximate molecular weights (kDa) are indicated on the left. A single asterisk (*) indicates the NRC4 resistosome, while a double asterisk (**) denotes the NRC4 dimer in the BN-PAGE data.

Analysis of the predicted model revealed that the pairwise distances between the four residues in the hydrophobic core are less than 6Å, indicating potential interactions among them (Fig. 2B). Furthermore, the hydrophobic core is predicted to be concealed by the α1 helix in the resting state (Fig. 2C). Interestingly, the predicted distances from L34 and A72 to L13 of α1, a critical residue in the MADA motif, are within 6Å, suggesting potential interaction between the hydrophobic core and the MADA motif (Fig. 2C). These observations imply that this hydrophobic core within the CC domain may play a role in NRC4 function by interacting with the MADA motif.

In the activated state, NRC4 forms a hexameric resistosome (Liu *et al*, 2024). Our AF3 structural analysis reveals that the identified hydrophobic core is embedded within the CC domain inside the activated resistosome complex (Fig. 2D). In addition, F44 and L128, the two residues not part of the hydrophobic core, are localized near the putative ion-conducting pore within the resistosome. Furthermore, one of the residues in the hydrophobic core, A72 of the α3 helix, is positioned just one residue upstream of the EDVID motif (Fig. S3B), which interacts with the R-cluster in the LRR domain to ensure proper NRC4 function (Liu *et al*, 2024). Collectively, our structural analysis suggests that this hydrophobic core in the NRC4 CC domain might play a role in maintaining the structural integrity necessary for intramolecular interactions among different functional motifs, thereby regulating NRC4 activation.

### The hydrophobic core is essential for NRC4 subcellular localization and membrane-associated puncta formation

NRC4 localizes in the cytoplasm in its resting state, and forms membrane-associated puncta upon activation (Duggan *et al*, 2021). To investigate whether the hydrophobic core contributes to NRC4 subcellular localization in both resting and activated states, we tagged NRC4 variants with a C-terminal GFP tag and performed cell biology experiments. Since wild-type NRC4 induces fast and robust cell death, limiting its application in cell biology studies, we employed the NRC4^L9A/V10A/L14A^ variant (NRC4^AAA^) that carries mutations in the MADA motif (Adachi *et al*, 2019a). This variant displays weaker and delayed cell death responses, allowing more accurate cell biology analysis (Fig. S5). We therefore used the NRC4^AAA^ variant as the background for subsequent functional analyses.

To investigate subcellular localization in the resting state, we co-expressed NRC4 variants with Rpi-blb2. We observed that the NRC4^AAA^ variant predominantly localized in the cytosol, consistent with previously reported localization of wild-type NRC4 (Fig. S6) (Duggan *et al*, 2021; Wang *et al*, 2023). Similarly, all four hydrophobic core mutants (L34E, A72E, I114E, and V118E) also predominantly localized in the cytosol at the resting state, indicating that mutations in the hydrophobic core do not affect NRC4 localization in the resting state (Fig. S6).

To determine whether the hydrophobic core affects NRC4 localization upon activation, we co-expressed the NRC4 mutants with Rpi-blb2 and its recognized effector AVRblb2. Upon activation, the NRC4^AAA^ variant exhibited a dramatic change in subcellular localization, forming membrane-associated puncta (Fig. 3A), consistent with the previous finding for the NRC4^L9E^ variant (Duggan *et al*, 2021). Interestingly, hydrophobic core mutants remained predominantly cytosolic when co-expressed with Rpi-blb2 and AVRblb2, suggesting that these mutations impair the ability of NRC4 to undergo activation-induced relocalization (Fig. 3A). However, the A72E mutant could still form a limited number of membrane-associated punctate structures upon Rpi-blb2 activation (Fig. 3A). We quantified cells with NRC4^AAA^ and NRC4^A72E^ puncta at 2 and 3 days post-infiltration (dpi) and found that NRC4^A72E^ exhibited delayed punctate formation (Fig. S7A). These findings suggest that NRC4^A72E^ has reduced efficiency and requires more time to oligomerize than NRC4^AAA^. Collectively, our results indicate that the hydrophobic core is essential for the efficient formation of NRC4 membrane-associated puncta upon activation of Rpi-blb2.

### Mutation of the hydrophobic core compromises NRC4 oligomerization

NRC4 oligomerizes into a hexameric resistosome upon activation (Contreras *et al*, 2023b; Liu *et al*, 2024). To determine whether mutations in the hydrophobic core affect NRC4 oligomerization, we conducted the Blue Native polyacrylamide gel electrophoresis (BN-PAGE) analysis using C-terminal Myc-tagged NRC4 variants. NRC4 and Rpi-blb2 were co-expressed with or without AVRblb2, and total proteins were collected for BN-PAGE analysis at 2 dpi. Consistent with previous reports using NRC4^L9E^, we observed that NRC4^AAA^ migrates as a ∼240 kDa complex at the resting state, and forms a ∼900 kDa complex at the activated state (Fig. 3B) (Contreras *et al*, 2023b). Importantly, we found that mutations in the hydrophobic core disrupted oligomerization, maintaining NRC4 in a dimeric state even in the presence of both AVRblb2 and Rpi-blb2 (Fig. 3B). Additionally, the NRC4^A72E^ mutant retained the ability to oligomerize at 3 dpi, forming a ∼900 kDa band similar to NRC4^AAA^ upon activation (Fig. S7B), consistent with its delayed puncta formation observed in microscopy. Collectively, these results demonstrate that the hydrophobic core plays a critical role in NRC4 oligomerization, providing a structural basis for its essential function in NRC4-mediated immunity.

### The hydrophobic core contributes to the NRC4-phospholipid association

A previous study demonstrated that five residues (K84, K87, K89, K91, and R94) within the disordered region of the NRC4 CC domain play a role in its association with phospholipids (Wang *et al*, 2023). To explore whether residues in the hydrophobic core also contribute to NRC4-phospholipid interactions, we performed phospholipid-binding assays using PIP strips. We conducted wheat-based *in vitro* translation to express the CC domain of NRC4^AAA^ and corresponding hydrophobic core mutants (Fig. 4A). After confirming protein accumulation, we incubated the proteins with PIP strips to determine the association between the NRC4 CC domain and various phospholipids. The fluorescent protein mCherry was used as a negative control to assess background binding.

**Figure 4.**
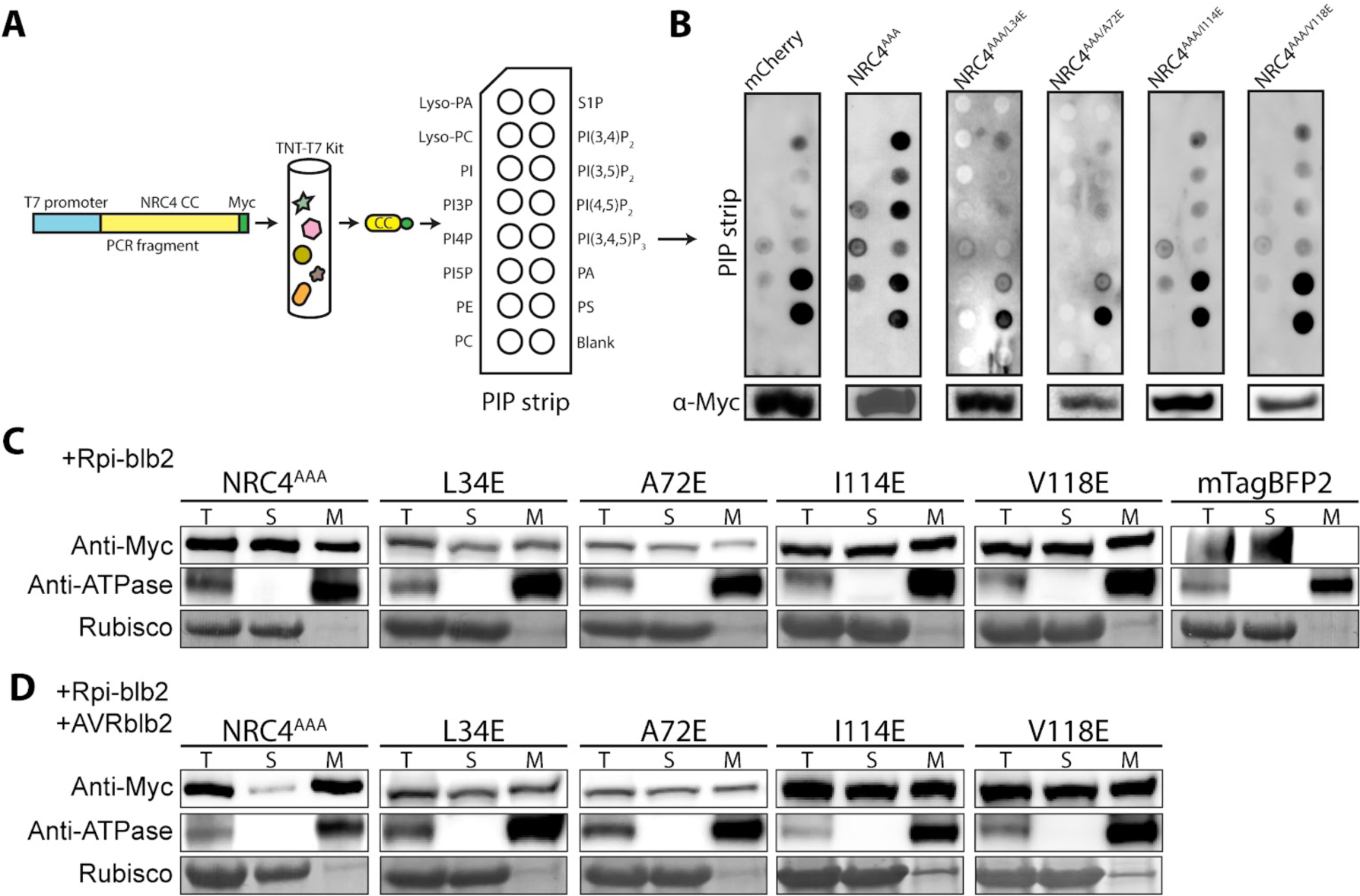
Mutations in the hydrophobic core compromise the NRC4 phospholipid association and membrane dynamics. (A) Experimental setup for investigating the association between the NRC4 CC domain and phospholipids. A gene fragment encoding the NRC4 CC domain with a C-terminal 4×Myc tag was amplified by PCR and *in vitro* translated using the TNT Coupled Wheat Germ Extract System. The resulting protein fractions were directly incubated with PIP strips for lipid association assays, and the NRC4 CC domain was detected using an anti-Myc antibody. (B) Phospholipid-binding assay for NRC4^AAA^ and hydrophobic core mutants. Myc-tagged mCherry served as a negative control. Expression levels of the *in vitro*-translated NRC4 CC domain variants were confirmed by western blot (shown below the binding assay results). (C) and (D) Membrane fractionation assay assessing the subcellular distribution of NRC4^AAA^ and hydrophobic core mutants co-expressed with Rpi-blb2, in the absence (C) or presence (D) of AVRblb2. Leaf samples were collected at 2 dpi, and total protein extracts were separated into soluble (S) and membrane (M) fractions. NRC4 proteins were detected using anti-Myc antibody. ATPase served as the membrane marker, while Rubisco is enriched in the soluble protein fraction. T represents the total extract. Rubisco was visualized by Coomassie blue staining.

We found that the NRC4^AAA^ CC domain is associated with PI4P and PI(4,5)P_2_, consistent with previous reports using the wild-type NRC4 CC domain (Fig. 4B) (Wang *et al*, 2023). Additionally, we observed the binding of NRC4^AAA^ to other phospholipids, including PI-monophosphates (PI3P, PI5P), PI-biphosphates [PI(3,4)P2, PI(3,5)P2], phosphatidic acid (PA) and phosphatidylserine (PS) (Fig. 4B). However, we noted that the negative control mCherry also binds to PA and PS, suggesting that signals detected with these lipids likely represent background binding in our assay.

In contrast, all variants harboring mutations in the hydrophobic core showed a substantially reduced ability to bind either the mono- or biphosphate forms of PI (Fig. 4B). All hydrophobic core mutants exhibited detectable protein accumulation levels, although variants A72E and I118E showed reduced accumulation (Fig. 4B). We next examined the lipid-binding ability of the two variants carrying mutations outside the hydrophobic core (F44E and L128E), both of which impaired NRC4-mediated cell death (Fig. S8). Interestingly, these mutations did not affect the interaction of the NRC4 CC domain with phospholipids (Fig. S8). Overall, our findings indicate that residues within the hydrophobic core are indispensable for NRC4-phospholipid association, whereas residues outside the core, such as F44 and L128, do not contribute to this association.

To further investigate the role of the hydrophobic core in membrane association, we conducted cell fractionation assays to assess how mutations in this region influence the membrane localization of full-length NRC4 under both resting conditions and upon activation. Under resting conditions, NRC4^AAA^ was detected in both soluble and membrane fractions at comparable levels (Fig. 4C). Upon activation with Rpi-blb2/AVRblb2, NRC4^AAA^ became significantly enriched in the membrane fraction, consistent with previous findings (Fig. 4D) (Duggan *et al*, 2021). However, in contrast to NRC4^AAA^, the hydrophobic core mutants failed to exhibit enhanced membrane association upon activation with Rpi-blb2/AVRblb2 (Fig. 4D). These results reveal that the hydrophobic core is important for NRC4-phospholipid association and NRC4 membrane enrichment, corresponding to the cell biology observations showing that the hydrophobic core of NRC4 is essential for its membrane-associated puncta formation and oligomerization upon activation by Rpi-blb2/AVRblb2 (Fig. 3).

### Disruption of the hydrophobic core does not affect NRC4 accumulation around the pathogen haustoria

NRC4 accumulates around the extrahaustorial membrane (EHM) of *P. infestans* during infection, likely facilitating effector detection (Duggan *et al*, 2021). To investigate whether residues in the hydrophobic core are important for NRC4 focal accumulation at the EHM, we expressed NRC4^AAA^ or its hydrophobic core mutants in *nrc2/3/4_KO N. benthamiana* leaves. At 1 dpi, we inoculated the detached leaves with *P. infestans* zoospores and examined the localization of NRC4 variants 2 days after pathogen inoculation. Consistent with previous reports on NRC4^WT^, NRC4^AAA^-GFP is focally accumulated at the EHM (Fig. 5) (Duggan *et al*, 2021). Similarly, all tested NRC4 hydrophobic core mutants displayed sharp signals around the EHM, but not in the cytosol or at the plasma membrane (Fig. 5). To further validate these observations, we utilized the NRC4^WT^-RFP as an EHM marker to validate the localization of the NRC4 mutants. Co-localization between the NRC4 hydrophobic core mutants and NRC4^WT^-RFP was observed, demonstrating that mutations in the hydrophobic core do not impair the capacity for EHM focal accumulation (Fig. S9). These findings reveal an interesting functional separation: while the hydrophobic core in the NRC4 CC domain is critical for resistosome formation, oligomerization, and phospholipid binding, it is dispensable for NRC4 focal accumulation at the EHM during pathogen infection. This suggests that distinct structural features within NRC4 mediate its different subcellular localization patterns during immune activation versus pathogen challenge.

**Figure 5.**
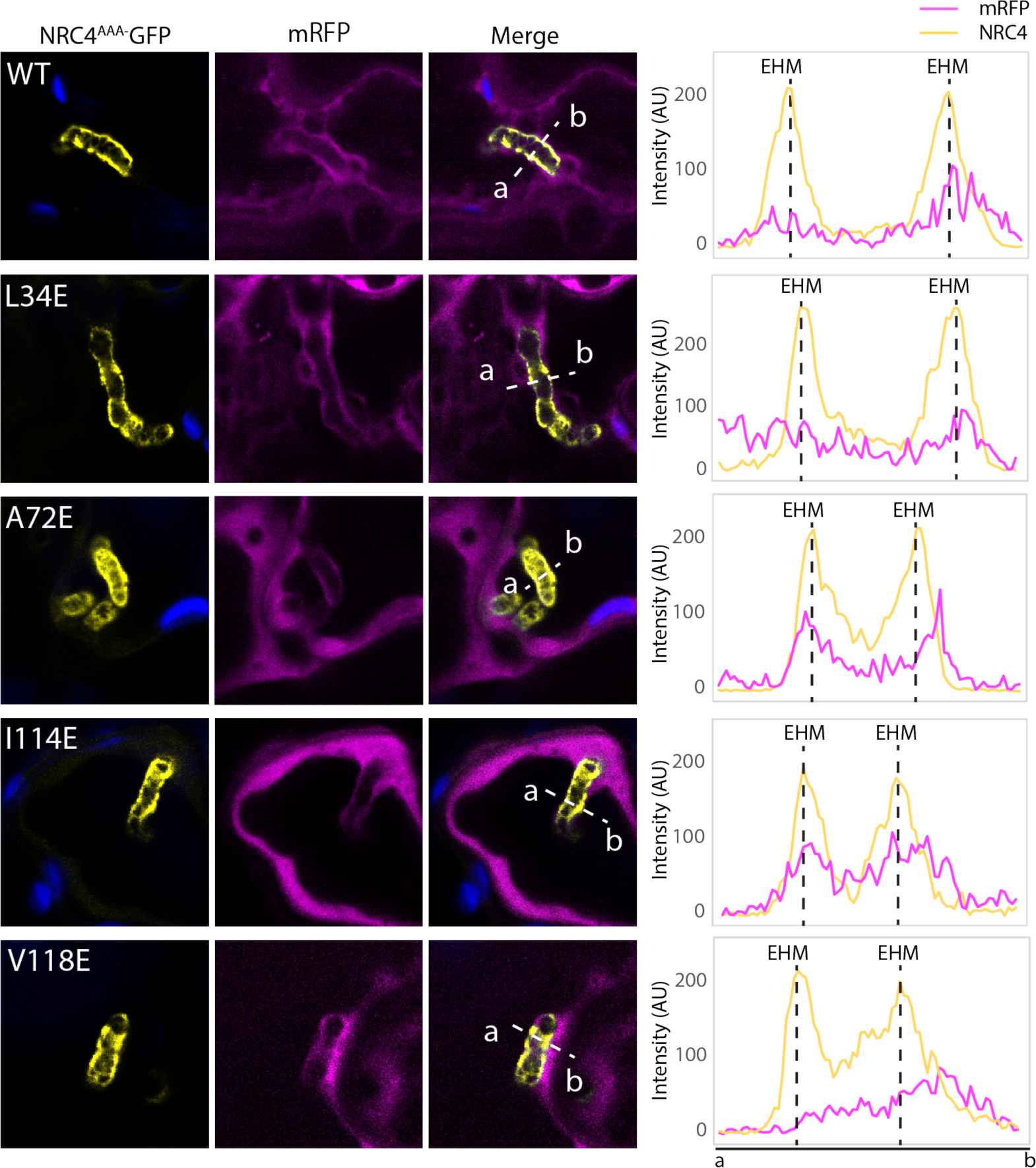
Mutations in the hydrophobic core do not affect NRC4 focal accumulation at the extrahaustorial membrane. Confocal images showing the accumulation of NRC4^AAA^ and hydrophobic core variants at the EHM. The NRC4 variants, tagged with C-terminal GFP, were transiently expressed with mRFP (cytosol marker) in *nrc2/3/4_KO N. benthamiana* leaves. At 1 dpi, *P. infestans* zoospores were inoculated onto detached leaves and incubated for two days. Images were taken at 3 dpi. The lines marked on the overlay panel indicated the selected region for measuring the intensity of each channel. The blue signal represents the chloroplast. Fluorescence intensity was measured from position “a” to “b” in each group, and the values are presented as arbitrary units (AU).

### The hydrophobic core within the CC domain is critical for the function of NRC helper NLRs and some singleton NLRs

Our discovery of a novel functional feature in the CC domain prompted us to investigate its conservation across other NLRs, focusing on those within the NRC superclade, as well as related singleton CNLs ZAR1, R2, and Rpi-vnt1 (Figure 6A). We selected 22 species representing diverse genera within the Solanaceae family from a recently published solanaceous NLRome dataset (Sugihara *et al*, 2023). From this dataset, we extracted 2,797 NLR sequences and grouped them into phylogenetic clades based on their NB-ARC domains (Fig. 6A, Fig. S10, File S1). Among these NLRs, a significant proportion of sensor NLRs in the NRC superclade contain N-terminal extensions that likely influence the canonical properties of the CC domain (Fig. S10) (Seong *et al*, 2020); therefore, we excluded these sensor NLRs from further analysis. From the remaining clades in the NRC superclade, we selected representative NLRs from each clade for detailed analysis: NRC2 from G8 clade, Rx from G12 clade, Bs2 from G2 clade, Rpi-amr1 from G14 clade, and Rpi-amr3 from G9 clade (Fig. 6A).

**Figure 6.**
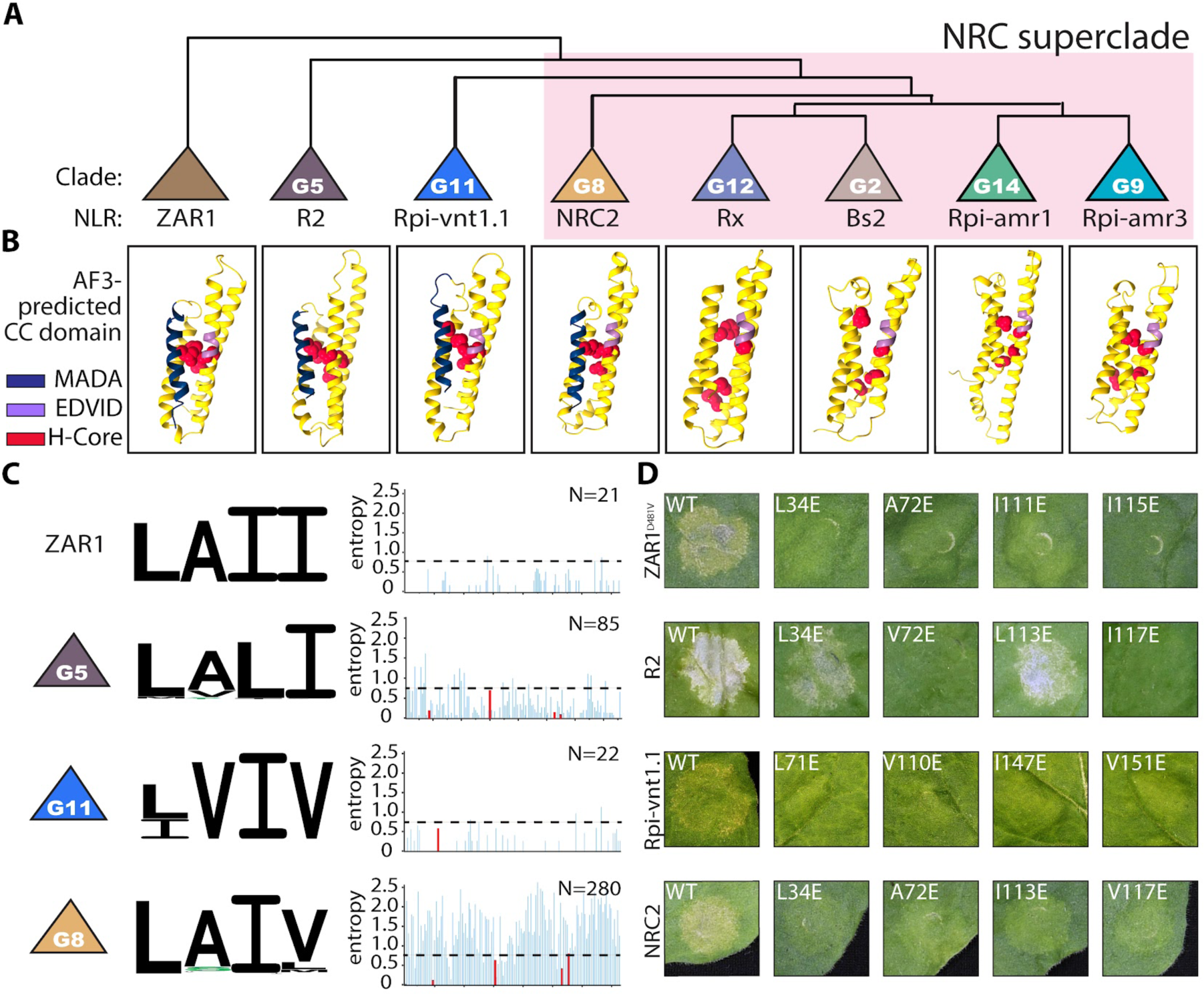
The hydrophobic core in the CC domain is essential for NRC2, ZAR1, and Rpi-vnt1.1. (A) Phylogenetic tree of the NRC superclade and related singleton NLRs, ZAR1, R2 and Rpi-vnt1.1. A representative NLR is indicated for each group. The NRC superclade is highlighted in pink. Sensor NLR clades possessing N-terminal extensions were excluded from this analysis. (B) AlphaFold3-predicted structures of selected NLRs. Red indicates hydrophobic core residues obtained from 2D alignment. H-Core represents the hydrophobic core. Blue indicates the MADA motif, and purple indicates the EDVID motif. (C) The amino acid composition of the hydrophobic core in each group is shown in the logo. Hydrophobic residues (AVLIPWFM) appear in black, hydrophilic residues (DE) in red, positively charged residues (KRH) in blue, and polar residues (STN) in green. From left to right, the positions correspond to NRC4 residues L34, A72, I114, and V118. Entropy values for the CC domain across singleton NLR ZAR1, R2, Rpi-vnt1.1, and NRC, with respective hydrophobic core residues in each clade highlighted in red. The number of NLR sequences in each clade is indicated in the top-right corner of the entropy panel. (D) Cell death phenotypes following hydrophobic core mutations in representative NLRs from each clade, photographed at 3 dpi. Quantitative data for these cell death assays can be found in Figure S11.

To determine whether the hydrophobic core feature is present in selected NLR groups, we aligned the CC domain sequences and identified the corresponding residues in each group (Files S2 to S9). These residues were then highlighted on the AlphaFold3-predicted CC domain structures of selected candidate NLRs. Fascinatingly, a compact hydrophobic core could be readily predicted in ZAR1, R2, Rpi-vnt1.1, and NRC2. In contrast, all representative NRC-dependent sensor NLRs lacked a predicted compact hydrophobic core (Fig. 6B). This pattern mirrors the absence of the MADA motif in these sensor NLRs, suggesting that during evolution, both the hydrophobic core and the MADA motif degenerated in NRC-dependent sensor NLRs. This finding further implies that the cell death-eliciting function in NLRs depends on both the MADA motif and the hydrophobic core.

Next, we performed sequence conservation analysis on the NLRs that form a hydrophobic core. The hydrophobic core residues in the G8 (NRC) clade were highly conserved, with most sequences containing LAIV (Fig. 6C). Entropy analysis revealed that these residues had entropy values lower than 0.75, signifying their relatively high sequence conservation (Fig. 6C). Similarly, the hydrophobic core residues in ZAR1, G5 clade, and G11 clade were also highly conserved, predominantly featuring LAII, LALI, and LVIV, respectively (Fig. 6C). Entropy analysis showed that the hydrophobic core residues in the G5 and G11 clades have low entropy values, with the ZAR1 clade showing no variation at these positions (Fig. 6C, Dataset S3, File S2).

To test whether the hydrophobic core is functionally conserved across these clades, we introduced point mutations in each of the residues in the hydrophobic core of representative NLRs from each clade. We first co-expressed wild-type NRC2 and its hydrophobic core mutants with Pto and effector AvrPto, which leads to Prf- and NRC2-dependent cell death, in *nrc2/3/4_KO* plants. All point mutations in the hydrophobic core of NRC2 compromised NRC2-mediated cell death, confirming the importance of the hydrophobic core in both NRC2 function (Fig. 6D, Fig. S11 A, B).

We then tested the functional importance of the hydrophobic core residues in ZAR1, R2 from G5 clade, and Rpi-vnt1 from G11 clade. For ZAR1 assays, we utilized the autoactive mutant ZAR1^D481V^ to elicit cell death. Cell death assays demonstrated that all hydrophobic core mutations in ZAR1^D481V^ significantly compromised its ability to mediate hypersensitive cell death, underscoring the critical role of the hydrophobic core in ZAR1 function (Fig. 6D, Fig. S11C, D). For Rpi-vnt1.1, we co-expressed Rpi-vnt1 or its hydrophobic core mutants with its cognate effector AVRvnt1 to elicit Rpi-vnt1.1-dependent cell death. Similar to ZAR1^D481V^, all the hydrophobic core mutants of Rpi-vnt1 compromised the cell death function of this NLR.

We then co-expressed wild-type R2 with its cognate effector AVR2 and observed strong hypersensitive cell death. Interestingly, while hydrophobic core mutants V72E and I117E significantly compromised R2 cell death function, the mutants L34E and L113E continued to exhibit strong cell death comparable to wild-type R2, suggesting that certain hydrophobic core residues in R2 may have functionally degenerated. Notably, R2 lacks the functional EDVID motif at the α3 helix (Fig. 6B, S12), unlike other tested candidate NLRs, implying that the presence of the EDVID motif is critical for the proper functional importance of the hydrophobic core in CNLs.

Altogether, our structural and functional analyses demonstrate that NRC helper NLRs and some singleton NLRs contain a compact hydrophobic core feature in the CC domain that is critical for NLR function. Additionally, the presence of both the MADA motif and the EDVID motif appears essential for the full functional significance of this hydrophobic core (Fig. 6B, D). As the hydrophobic core is strategically positioned between these two important functional motifs, we propose that these structural elements cooperate to execute the hypersensitive cell death-eliciting function of NLRs.

## Discussion

While recent cryo-EM studies unveiled detailed structures of NRC complexes in both resting and activated states, the precise structural features of the CC domain were not fully resolved, likely due to the high flexibility of the region connecting the CC and NB-ARC domains (Liu *et al*, 2024; Madhuprakash *et al*, 2024; Selvaraj *et al*, 2024). Building upon the previously established knowledge of the MADA motif in the α1 helix, our study identified several conserved hydrophobic residues across NRCs within the α2 to α4 helices (Fig. S1). We further demonstrated that several of these hydrophobic residues, situated at the core of the CC domain, form a hydrophobic core essential for NRC4 function (Fig. 1 and 2).

Several CNL-type singleton and helper NLRs undergo dynamic subcellular transitions from the cytosol to the plasma membrane, where they form pentameric or hexameric resistosome complexes upon activation (Wang *et al*, 2019a; Bi *et al*, 2021; Duggan *et al*, 2021; Liu *et al*, 2024). We found that mutations in the hydrophobic core do not affect the subcellular dynamics of NRC4 in its resting state (Fig. S6). However, upon activation of the sensor NLR Rpi-blb2, these mutations hinder the ability of NRC4 to oligomerize and localize to the plasma membrane (Fig. 3A). Additionally, BN-PAGE analysis showed that the hydrophobic core mutants remain as dimers when co-expressed with Rpi-blb2 and AVRblb2 (Fig. 3B). One possible explanation is that NRC4 hydrophobic core mutants are unable to receive activation signals from sensor NLRs, remaining as inactive homodimers in the cytosol. Alternatively, the hydrophobic core mutant NRC4 can receive the signal, but the mutation altered the CC domain structure, preventing the disruption of the homodimer and the formation of resistosome complexes.

While the MADA motif is an essential feature in the α1 helix of NRC4, the detailed mechanism of how hydrophobic residues in the MADA motif contribute to NRC4 function remains unclear (Adachi *et al*, 2019a). Using AF3 predictions, we showed that the L13 residue within the MADA motif, which is critical for NRC4 cell death activity, potentially interacts with two hydrophobic core residues in the α2 and α3 helices (Fig. 2C). The ZAR1 resistosome study proposed a “flipping out” process of the α1 helix in the CC domain during ZAR1 activation, a crucial event for ZAR1-mediated cell death (Wang *et al*, 2019a). Prior structural analysis of NRCs failed to identify the α1 helix of the CC domain in the activated resistosome, suggesting that this helix might also undergo the “flipping out” process (Liu *et al*, 2024; Madhuprakash *et al*, 2024). Given the similarity between ZAR1 and NRC resistosomes, we speculate that mutations in this hydrophobic core could disrupt interactions between the MADA motif and the α2-α3 helices, impairing the “flipping-out” process of the α1 helix during activation and ultimately compromising ZAR1 and NRC function.

A recent cryo-EM study showed that a triple mutation within the EDVID motif of the α3 helix disrupts NRC4-mediated cell death, while single mutations in this region do not affect NRC4 functionality (Liu *et al*, 2024). In contrast, we observed that a single mutation at A72, located adjacent to the conserved EDVID motif in the hydrophobic core, is sufficient to impair NRC4-mediated cell death (Fig. 1C). Interestingly, NRC4 with the A72 mutation can still oligomerize and localize to the plasma membrane at 3 dpi. However, the A72E mutant resistosome is non-functional and unable to trigger cell death. Our findings suggest that A72 likely plays a role in modulating interactions between the EDVID motif and the R-cluster, thus regulating NRC4-mediated cell death (Fig. 1C, Fig. S3B).

The interaction between phospholipids and helper NLRs is crucial for NLR-mediated cell death (Saile *et al*, 2021; Wang *et al*, 2023). NRC4 has previously been shown to associate with PI4P and PI(4,5)P_2_, two key phospholipids located on the plasma membrane (Wang *et al*, 2023). A positively charged cluster in the α4 helix of NRC4, conserved among other helper NLRs such as ADR1 and NRG1, plays a vital role in NRC4-phospholipid interaction (Wang *et al*, 2023). In our study, single mutations in the hydrophobic core of NRC4 disrupted its association with phospholipids (Fig. 4B). Since the hydrophobic core is embedded within the resistosome, it is unlikely to interact directly with lipids in the activated state (Fig. 2D). A possible explanation is that these mutations alter the structure of the NRC4 CC domain, disrupting the proper conformation of the positively charged loop and thereby impairing interactions between NRC4 and phospholipids. Notably, most hydrophobic core mutants of NRC4 lose both their capacity to associate with phospholipids and their ability to oligomerize. Our results support the hypothesis that NRC4-phospholipid interactions are crucial for resistosome formation. When the CC domain structure is disrupted, NRC4-phospholipid binding is compromised, causing NRC4 to remain in a cytosolic homodimer state rather than oligomerizing into functional resistosome complexes (Fig. 3 and 4).

The focal accumulation of NRC4 on the EHM is a distinctive feature observed during *P. infestans* infection (Duggan *et al*, 2021). Interestingly, NRC4 hydrophobic core mutants can still focally accumulate on the EHM during infection, indicating that these mutations do not affect all intrinsic features of NRC4. Previous studies on phospholipid composition at the EHM identified PI4P and PI(4,5)P_2_ as the primary phospholipids enriched at the EHM (Rausche *et al*, 2021). However, our lipid-binding assay reveals that hydrophobic core mutations of NRC4 abolish the ability to associate with both PI4P and PI(4,5)P_2_ (Fig. 4B). One possible explanation is that the focal accumulation of NRC4 on the EHM may not depend on direct association of the NRC4 CC domain with lipids. Instead, other regions of the NRC4 canonical domain or interactions with host or pathogen proteins are likely involved in this process. NRC4 may require these additional domains or partnering proteins to facilitate NRC4 focal accumulation on the EHM.

Previous studies suggest that NRC helper NLRs and NRC-dependent sensor NLRs arose through sub-functionalization from an ancestral singleton NLR (Contreras *et al*, 2023a). This process likely involved divergence of sequence features, resulting in altered protein functionality. One notable example is the MADA motif, which is conserved among NRCs, ZAR1, and some sensor NLRs, but not in NRC-dependent sensor NLRs (Adachi *et al*, 2019a). Similarly, while the hydrophobic core described here is conserved in some singleton CNLs and NRC helper NLRs, it is missing in NRC-dependent sensor NLRs (Fig. 6B). We propose that together with the MADA motif, the hydrophobic core emerged in the common ancestral NLR of singleton CNLs and NRC helper NLRs, but subsequently degenerated in NRC-dependent sensor NLRs.

Although the hydrophobic core is present in R2, only two of the mutants demonstrated reduced cell death (Fig. 6D). In both ZAR1 and NRCs, activation triggers the formation of pentameric or hexameric resistosomes, which initiate downstream hypersensitive responses (Wang *et al*, 2019a; Madhuprakash *et al*, 2024; Liu *et al*, 2024). However, it remains unclear whether R2 forms a resistosome similar to those of NRCs or ZAR1 and induces cell death through a comparable mechanism. Notably, R2 contains the MADA motif, which is critical for cell death initiation in many NLRs. In contrast, the EDVID motif—a conserved feature found in ZAR1, Rpi-vnt1.1, NRCs, and many other NLRs—is absent in R2 (Fig. 6B, S12) (Förderer *et al*, 2022; Rairdan *et al*, 2008; Wróblewski *et al*, 2018). The presence or absence of MADA and EDVID motifs, coupled with the variable requirement for the hydrophobic core described here, suggests that different types of CNLs possess unique structural and sequence features within their CC domains, resulting in distinct regulatory mechanisms (Fig. 6).

In conclusion, our study identifies a previously uncharacterized hydrophobic core in the CC domain of NRC helper NLRs and some singleton NLRs that is essential for their functions in immune signaling. This hydrophobic core contributes to resistosome formation, phospholipid binding, and subcellular dynamics, but is dispensable for NRC4 focal accumulation at the EHM during pathogen infection. Furthermore, our analysis reveals that this structural feature is highly conserved in NRC helper NLRs and some singleton NLRs but has diverged in sensor NLRs, providing new insights into the structure-function connection of NLR-mediated immunity in plants.

## Materials and Methods

### Plant material and growth conditions

Wild-type and *nrc2/3/4_KO N. benthamiana* plants were grown in a walk-in growth chamber maintained at 25L°C, under a 16 h light/8 h dark cycle (Wu *et al*, 2020).

### Plasmid construction

The CC, NB-ARC, and LRR domains of NRC2/4 were modularized and cloned into the Golden Gate Level 0 acceptor vector pAGM9121 (Weber *et al*, 2011). Site-directed mutagenesis were performed on the CC domain Level 0 module using inverse PCR, and then digested with AarI (NEB) and ligated with T4 DNA ligase (Thermo Scientific). The mutated CC domain modules were assembled with the NB-ARC and LRR modules, along with a C-terminal 4x Myc tag or GFP tag, into the pICH86988 binary vector, containing the 35S promoter and OCS terminator, using Golden Gate assembly (Weber *et al*, 2011; Engler *et al*, 2014). The pICH86988 constructs containing NRC4-myc or GFP were subsequently transformed into *Agrobacterium tumefaciens* strain GV3101(pMP90). For *in vitro* translation experiments, the CC domain module of NRC4 (wild-type and hydrophobic core mutants) was assembled with the T7 promoter, CMV1 5’ UTR, T7 terminator, and an overhang-modified C-terminal Myc tag (CTCA-GCTT) into the acceptor vector pICSL86900OD (Engler *et al*, 2014). To study the hydrophobic core function in R2 and Rpi-vnt1.1, the CDS of R2 and Rpi-vnt1.1 were cloned into Level 0 acceptor pAGM9121(Tai *et al*, 1999; Foster *et al*, 2009). For ZAR1, a D481V mutation variant (autoactive) Level 0 plasmid was used (Contreras *et al*, 2024). Site-directed mutagenesis in the CC domain was performed as described above. The resulting NLR variants were subcloned into pICH86988 for cell death assays. The effector gene AVR2 was synthesized and assembled into pICH47751 vectors, incorporating the 35S promoter and OCS terminator (Gilroy *et al*, 2007; Lin *et al*, 2022). The primers used for cloning and the plasmids used in this study are listed in Datasets S1 and S2.

### Agroinfiltration

*A. tumefaciens* GV3101(pMP90) was used to express NLRs and effectors. The *Agrobacterium* stocks were refreshed on 523 medium containing the appropriate antibiotics and incubated overnight at 28°C. The cells were harvested by centrifugation at 5000 × g for 5 minutes and resuspended in MMA buffer (10 mM MgCl_2_, 10 mM MES-KOH, 150 μM acetosyringone, pH 5.6). The cell concentrations were adjusted to 0.2 for all NLRs and 0.1 for all effectors for subsequent experiments. *Agrobacterium* suspensions with the desired optical density (OD) were infiltrated into *N. benthamiana* leaves using a 1 mL syringe.

### Cell death assay

The complementation assays were performed in the *nrc2/3/4*_KO *N. benthamiana*, while functional analyses of hydrophobic core-mutated R2, ZAR1, and Rpi-vnt1.1 were conducted in wild-type *N. benthamiana*. The NLR proteins and their corresponding effectors were transiently expressed in *N. benthamiana* using *A. tumefaciens* strains carrying expression vectors for the indicated proteins. Cell death intensity was quantified using the UVP ChemStudio Imaging System (Analytik Jena) at 6 dpi for NRC4 mutants screen and at 3 dpi for R2, ZAR1, NRC2/3, and Rpi-vnt1.1 mutants. Autofluorescence of necrotic regions was detected using the blue LED excitation light and a FITC emission filter (519 nm). Cell death intensity was normalized by dividing the pixel value by the maximum signal pixel value (65535) detected by the UVP ChemStudio.

### Protein extraction for SDS-PAGE analysis

The protein extraction method was conducted as previously described (Win *et al*, 2011). The leaf disc was homogenized with ceramic beads in a tissue homogenizer under liquid nitrogen. Total protein was extracted with GTEN buffer comprising 10% glycerol, 25 mM Tris (pH 7.5), 1 mM EDTA, 150 mM NaCl, 2% (w/v) PVPP, 10 mM DTT, 1× protease inhibitor cocktail (Sigma, P9599), and 0.2% IGEPAL (Sigma). The protein extract was centrifuged at 5000 g for 10 minutes at 4°C, and the supernatant was transferred to a new Eppendorf tube and centrifuged again at 13000 ×g for 5 minutes at 4 °C to remove the remaining debris as possible. The final supernatant was combined with 4x sample loading dye containing 200 mM Tris-HCl (pH 6.8), 8% (w/v) SDS, 40% (v/v) glycerol, 50 mM EDTA, 0.08% bromophenol blue, and 0.1 M DTT. The protein samples were incubated at 70 °C for 10 minutes before SDS-PAGE analysis.

### SDS-PAGE assay and immunodetection analysis

SDS-PAGE electrophoresis and immunodetection analyses were conducted as previously described (Win *et al*, 2011). Denatured samples were separated on PAGE gels prepared with 10% or 15% T-Pro EZ gel solution (Omics bio), containing 0.1% TEMED (Bio-Rad) and 0.1% ammonium persulfate (Bio-Rad). Proteins were transferred to a PVDF membrane using the Trans-Blot Turbo Transfer System (Bio-Rad). Following the transfer, the membrane was washed with Tris buffer saline (TBS) with 0.1% Tween-20 and blocked with 5% nonfat dry milk. For immunodetection, anti-Myc (GenScript, A00704) and anti-H+ATPase (Agrisera, AS07 260) were used as primary antibodies for NRC4 protein and membrane fraction detection, respectively. Secondary antibodies included Peroxidase AffiniPure Goat Anti-Mouse IgG (H+L) (Jackson, 115-035-003) and Peroxidase Conjugated Goat Anti-Rabbit IgG (H+L) (Sigma, AP132P). Primary antibodies were diluted at 1:5000, and the secondary antibodies were diluted at 1:20000. Chemiluminescence signals were detected using a mixture of SuperSignal West Pico PLUS Chemiluminescent Substrate (Thermo Scientific, 34580) and SuperSignal West Femto Maximum Sensitivity Substrate (Thermo Scientific, 34096) in a 3:1 ratio. Images were captured using the UVP ChemStudio Imaging System. SimplyBlue SafeStain (Invitrogen, 465034) was used to visualize the Rubisco signal on the PVDF membrane.

### Blue Native PAGE

The BN-PAGE method was conducted as previously described (Contreras *et al*, 2023b). To assess whether oligomerization was affected in the mutants, we transiently expressed the NRC4 hydrophobic core mutants along with Rpi-blb2, with or without AVRblb2, in 4-week-old plants. Leaf discs were collected at 2 and 3 dpi, and total protein was extracted using a native PAGE extraction buffer containing 50 mM HEPES (pH 7.5), 50 mM NaCl, 5 mM MgCl2, 10% glycerol, 10 mM DTT, 1x protease inhibitor cocktail (Sigma, P9599), and 1% digitonin (Sigma, D141). Protein extracts were incubated on ice at 4 °C for 10 minutes, followed by centrifugation at 12,000 rpm for 15 minutes at 4 °C. The supernatant was transferred to a new tube and centrifuged again at 12,000 rpm for 10 minutes at 4 °C to avoid the interference of the pellet. The resulting protein extracts were diluted to 0.4x using the native PAGE extraction buffer. Diluted samples were then mixed with 4x sample buffer and NativePAGE 5% G-250 Sample Additive to achieve a final G-250 concentration of 0.125%. Samples were loaded onto a 4–16% Native PAGE gel (Invitrogen, BN1002) and electrophoresis was carried out following the manufacturer’s protocol. In brief, Native PAGE was performed using the XCell system (Invitrogen) at 4 °C. The gel was run with a dark cathode buffer at 150 V for 45 minutes, followed by replacing the dark cathode buffer with a light cathode buffer and continuing at 200 V for 120 minutes. After electrophoresis, proteins were transferred to a PVDF membrane using the Trans-Blot Turbo Transfer System (Bio-Rad). The membrane was fixed with 8% acetic acid, air-dried in a chemical hood, and subsequently rinsed with 99% ethanol. The membrane was washed with ddH_2_O and TBST to remove residual ethanol before the immunodetection mentioned above.

### *In vitro* translation

The plasmid containing the *T7-CMV1:NRC4 CC-Myc* fragment was amplified using the T7 promoter forward primer and the T7 terminator reverse primer with the KAPA HiFi HotStart ReadyMix (Roche). The resulting PCR fragments were purified using a FavorPrep GEL/PCR purification Mini kit (Favorgen). The *in vitro* translation assay was performed following the protocol of the TNT-T7 coupled wheat germ extract system (Promega). Briefly, 1 µg of purified PCR fragments was mixed with the TNT wheat germ extract, TNT reaction buffer, amino acid mixture, RNase inhibitor, and T7 polymerase. The reaction mixture was incubated at 30°C for 2 hours. The NRC4-CC domain tagged with 4× Myc was subsequently analyzed by immunoblotting as mentioned above.

### Lipid binding assay

The PIP strips (Invitrogen) were first blocked by incubating with 3% bovine serum albumin (Fraction V, low heavy metal, MERK) at room temperature for one hour. The NRC4 CC domain proteins, obtained from *in vitro* translation, were then incubated with the PIP strips at 4°C overnight. Following incubation, the PIP strips were washed with TBST before the immunodetection assay as described above.

### Protein fractionation

Membrane enrichment was performed using a slightly modified version of a previously described protocol (Abas & Luschnig, 2010). NRC4 was transiently co-expressed with *Rpi-blb2*, with or without *AVRblb2*, in *nrc2/3/4_KO N. benthamiana*. At 2 dpi, leaves were harvested and homogenized under liquid nitrogen using a pestle and mortar. The powdered leaf tissue was mixed with membrane protein extraction buffer (0.33 M sucrose, 20 mM Tris-HCl [pH 8], 1 mM EDTA), supplemented with 5 mM DTT, 1× protease inhibitor (Thermo Scientific), and 0.5% PVPP. The mixture was first centrifuged at 2000 ×g, and the supernatants were further filtered through Miracloth (MERK, 475855-1R). The supernatant was transferred to a new Eppendorf tube, and 50 µl was set aside as the total protein fraction. The remaining supernatant was ultracentrifuged at 21,000 ×g for 1.5 hours to separate the soluble protein fraction and the membrane protein. The soluble protein fraction was transferred to a new Eppendorf tube, while the pellet was resuspended in a membrane extraction buffer (without PVPP) at one-third the volume of the soluble protein fraction. The membrane fraction was then diluted threefold with a membrane extraction buffer (without PVPP) before being applied to SDS-PAGE and immunoblotting analysis.

### Confocal microscopy

To analyze the subcellular localization of wild-type and hydrophobic core-mutated NRC4, four-week-old *nrc2/3/4_KO N. benthamiana* plants were agroinfiltrated with NRC4-GFP, HF-Rpi-blb2, mTagBFP, SOBIR1-mCherry, and either with or without Flag-AVRblb2. At 2 or 3 dpi, discs of the infiltrated leaves were imaged using an Andor Dragonfly spinning disc confocal microscope (Andor) equipped with a 40× water immersion objective lens (Advanced optical microscope core facility, Academia Sinica) or Leica Stellaris 8 inverted confocal microscope with 40x water immersion objective lens (Live-cell-image core lab, Institute of Plant and Microbiology, Academia Sinica). Excitation wavelengths of 405 nm, 488 nm, and 561 nm were used for BFP, GFP, and mCherry, respectively. The corresponding emission wavelengths were 422–468 nm for BFP, 505–550 nm for GFP, and 572–615 nm for mCherry. Chloroplast autofluorescence was labeled using excitation at 637 nm and emission in the range of 660–737 nm. For imaging of NRC4 haustoria accumulation, *P. infestans* inoculation was performed as described below. After 1 day of inoculation, leaf discs from the infected regions were imaged using a 40x water immersion objective lens on a Leica Stellaris 8 inverted confocal microscope as described before (Yuen *et al*, 2024a). The single-plane images were exported to Fiji (ImageJ) software for quantitative analysis. GFP and RFP signals were measured at selected areas (5 µm) spanning the peripheral cytoplasm and across the extrahaustorial membrane (EHM).

### Inoculation of Phytophthora infestans

*P. infestans* 214009 (a gift from Dr. Jin-Hsing Huang, Taiwan Agriculture Research Institute) was originally isolated from Taiping, Taichung City, Taiwan. *P. infestans* was maintained on Rye sucrose medium at 19°C under dark conditions. Before performing the focal accumulation assay for NRC4, *P. infestans* was subcultured on detached *N. benthamiana* leaves. For the EHM focal accumulation assay, leaves from 4-week-old *nrc2/3/4*_KO *N. benthamiana* plants were infiltrated with *Agrobacterium* containing either wild-type or hydrophobic core mutant NRC4 constructs and a cytosolic RFP marker. Zoospores were collected from sporangia on *P. infestans*-infected *N. benthamiana* leaves by incubating the sporangia in sterile water at 4°C as previously described (Yuen *et al*, 2024b). The zoospores were then inoculated onto detached *N. benthamiana* leaves 2 days after transient expression of NRC4 variants.

### Protein structure prediction

To identify the position of the conserved hydrophobic residue candidates, we applied AlphaFold 3 to predict the NRC4 monomer (pTM=0.82) (Abramson *et al*, 2024). Assuming similar protein conformation, we superimposed the NRC4 monomer to the published NbNRC2 dimer (PDB: 8RFH) by using the “Matchmaker” function to further correct the NbNRC4 structure we obtained from AlphaFold3 (Abramson *et al*, 2024). Since the CC domain is missing in this NbNRC2 homodimer structure and may make the CC domain differ from the current model, we cross-compared the position of the hydrophobic core on the NbNRC4 resistosome structure (PDB: 9CC8).

### Phylogenetic analysis

The amino acid sequences of the NB-ARC domain of the NRC superfamily from 20 species representing various genera within the Solanaceae were extracted from the dataset in the NLRdome (Sugihara *et al*, 2023). A total of 2,581 sequences, including reference NLR sequences, were aligned using MAFFT (Katoh *et al*, 2019). Gaps in the aligned sequences were manually removed in MEGA (Molecular Evolutionary Genetics Analysis), and the trimmed amino acid sequences were used for phylogenetic analysis in IQtree (Kumar *et al*, 2016; Minh *et al*, 2020). The phylogenetic analysis employed the Maximum Likelihood method with the JTT+F+R10 evolutionary model and 1000 bootstrap tests to assess the reliability of the tree. NLR clades were further categorized based on the grouping criteria outlined in the previous study (Seong *et al*, 2020).

### Hydrophobic core amino acid logo creation and entropy analysis

The full-length NLR amino acid sequences and the CC domain sequences from each clade were extracted and aligned using MAFFT, with NRC2/3/4 serving as the reference for identifying the hydrophobic core. To create the amino acid logo, the corresponding residues of the hydrophobic core in each clade were extracted, and the reference NRC2/3/4 sequences were removed. Amino acid logos for each clade were then generated based on the extracted hydrophobic core residues using WebLogo 3 (https://weblogo.threeplusone.com/create.cgi) (Crooks *et al*, 2004). After CC domain alignment and removal of NRC2/3/4 sequences, the remaining sequences were subjected to entropy analysis. Shannon entropy was calculated using an online tool developed by the Los Alamos National Laboratory (https://www.hiv.lanl.gov/content/sequence/ENTROPY/entropy.html). The hydrophobic core was labeled, and a cutoff score of 0.75 was applied to identify highly variable residues compared to the NRC group. Detailed analysis results are included in the Dataset S3.

## Acknowledgments

We thank Mark Youles (SynBio, The Sainsbury Laboratory, UK) for sharing plasmids for molecular cloning. We thank Mei-Jane Fang and Ji-Ying Huang in the Live-Cell-Imaging Core Lab (Institute of Plant and Microbial Biology, Academia Sinica, Taiwan) and Shu-Chen Shen in the Advanced Optical Microscope Core Facility (Agricultural Biotechnology Research Center, Academia Sinica, Taiwan) for help with confocal imaging. We thank Dr. Jin-Hsing Huang (Taiwan Agriculture Research Institute) for providing the *P. infestans* isolate 214009 and advice on pathogen inoculation. This project was funded by the 2030 Emerging Young Scholar Program of the National Science and Technology Council (NSTC) of Taiwan (NSTC-112-2628-B-001-007), a bilateral exchange grant of NSTC and Royal Society (UK) (NSTC-113-2927-I-001-514, IEC\NSFC\233289), and the intramural fund of Institute of Plant and Microbial Biology, Academia Sinica, Taiwan. ELHY and TB are funded by Biotechnology and Biological Sciences Research Council (BBSRC) BB/X016382/1.

## Author Contributions

HYW and CHW contributed to the conceptualization of the project. HYW contributed to data curation. HYW, ELHY, and CHW contributed to the formal analysis. HWY, ELHY, and CHW contributed to the investigation. KTL and FJG contributed to the methodology. TOB and CHW supervised the research and acquired funding. CHW managed the project. HYW and ELHY contributed to data visualization. HYW and CHW wrote the initial manuscript draft. HYW, ELHY, and CHW edited the manuscript.

## Data Availability

All study data are included in the article and/or supporting information.

## Supplementary Data

Figure S1. Alignment of the CC domain of NbNRC2/3/4 and AtZAR1.

Figure S2. The protein accumulation of NRC4 hydrophobic residues mutant variants.

Figure S3. The localization of the hydrophobic core in the NRC4 dimer and the resistosome.

Figure S4. The hydrophobic core of the CC domain in the NRC2 dimer and the resistosome

Figure S5. NRC4^AAA^ showed no cell death at 2 dpi compared to wild-type NRC4.

Figure S6. Mutations on the hydrophobic core do not affect NRC4 subcellular localization in the resting state.

Figure S7. The mutation on A72 delays the formation of the resistosome and puncta on the plasma membrane.

Figure S8. Mutations of hydrophobic residues outside the hydrophobic core did not affect the association of the NRC4 CC domain with phospholipids.

Figure S9. Mutations of the hydrophobic core do not affect the NRC4 focal accumulation on the EHM.

Figure S10. Phylogenetic tree of the NRC superclade and selected singleton NLR groups (R2, Rpi-vnt1, and ZAR1).

Figure S11. Quantitative cell death result of NRC2, NRC3, R2, and ZAR1 variants.

Figure S12. The EDVID motif is absent in R2.

Table S1. List of the constructs used in cell death assays.

Table S2. List of the constructs used in lipid binding assays.

Table S3. List of the constructs used in cell biology assays.

Table S4. List of the constructs used in BN-PAGE assays.

Table S5. List of the constructs used in cell fractionation assays.

Table S6. List of the constructs used in focal assays.

Dataset S1. List of primers used in this study. (XLSX)

Dataset S2. List of plasmids used in this study. (XLSX)

Dataset S3. Results of entropy analysis. (XLSX)

File S1. NRC superfamily phylogenetic tree files of 22 species.

File S2. ZAR1 group CC domain alignment. (TXT)

File S3. G5 group CC domain alignment. (TXT)

File S4. G11 group CC domain alignment. (TXT)

File S5. G8 group CC domain alignment. (TXT)

File S6. G12 group CC domain alignment. (TXT)

File S7. G2 group CC domain alignment. (TXT)

File S8. G9 group CC domain alignment. (TXT)

File S9. G14 group CC domain alignment. (TXT)

