## Supplementary material for "A hydrophobic core in the coiled-coil domain is essential for NRC resistosome function": Supplementary Figures and Tables.pdf

**This document include:**

Supplementary Figures S1 to S12

Tables S1 to S6

Appendix References

**Other Materials for this manuscript include the following (not in this Appendix file):**

Dataset S1. List of primers used in this study. (XLSX)

Dataset S2. List of plasmids used in this study. (XLSX)

Dataset S3. Results of entropy analysis. (XLSX)

File S1. NRC superfamily phylogenetic tree files of 22 species.

File S2. ZAR1 group CC domain alignment. (TXT)

File S3. G5 group CC domain alignment. (TXT)

File S4. G11 group CC domain alignment. (TXT)

File S5. G8 group CC domain alignment. (TXT)

File S6. G12 group CC domain alignment. (TXT)

File S7. G2 group CC domain alignment. (TXT)

File S8. G9 group CC domain alignment. (TXT)

File S9. G14 group CC domain alignment. (TXT)

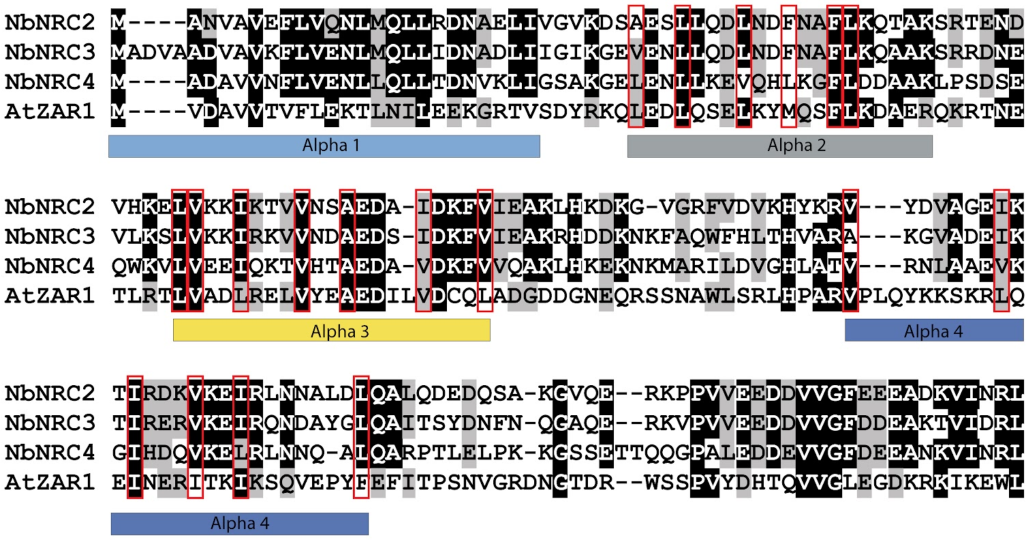


**Figure S1. Alignment of the CC domain of NbNRC2/3/4 and AtZAR1.** Conserved hydrophobic residues within α2–α4 are indicated by red rectangles. Amino acids highlighted with a black background are identical in at least three reference sequences, while those with a gray background are similar but not identical in at least three reference sequences.

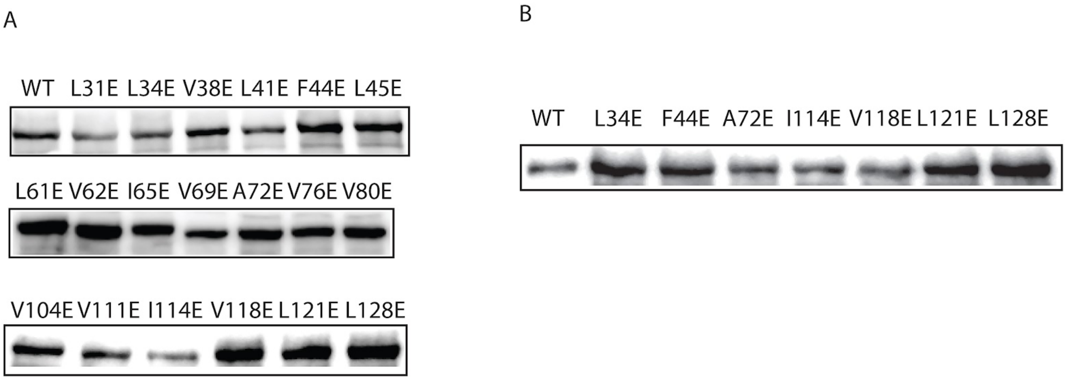


**Figure S2. The protein accumulation of NRC4 hydrophobic residues mutant variants.** (A) Protein accumulation levels of NRC4 hydrophobic residue mutants in the resting state. (B) Protein accumulation levels of NRC4 hydrophobic residue mutants that diminish NRC4-mediated cell death following Rpi-blb2 activation. Wild-type NRC4 and its hydrophobic residue mutants were tagged at the C-terminus with Myc, and protein accumulation was evaluated by immunoblotting using an anti-Myc antibody.

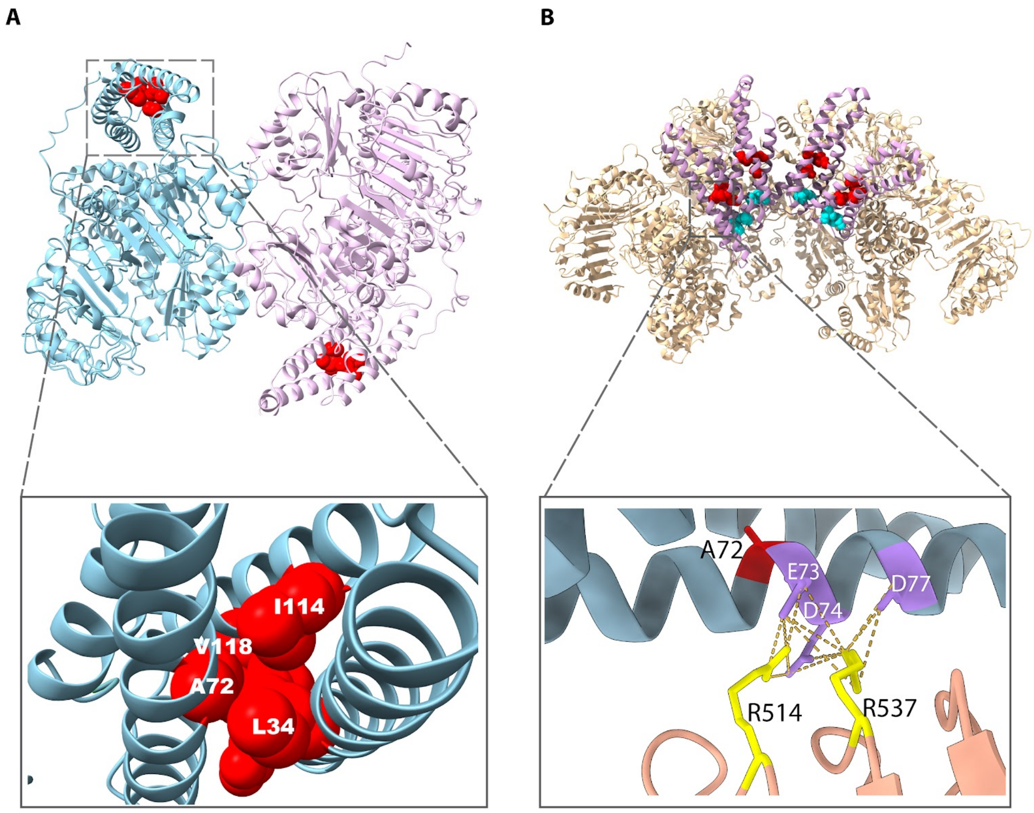


**Figure S3. The localization of the hydrophobic core in the NRC4 dimer and the resistosome.** Residues L34, A72, I114, and V118 are centrally located in the NRC4 CC domain, forming a hydrophobic core. The illustrated dimer is the predicted NbNRC4 structure based on the NbNRC2 dimer (PDB: 8RFH). The lower panel is the side view of the CC domain, highlighting the hydrophobic core residues in red. (B) The localization of EDVID and R-Cluster on the NRC4 resistosome structure (PDB=9CC8). Lower panel: The position of A72 (Red), EDVID motif (Purple), and R cluster (Yellow).

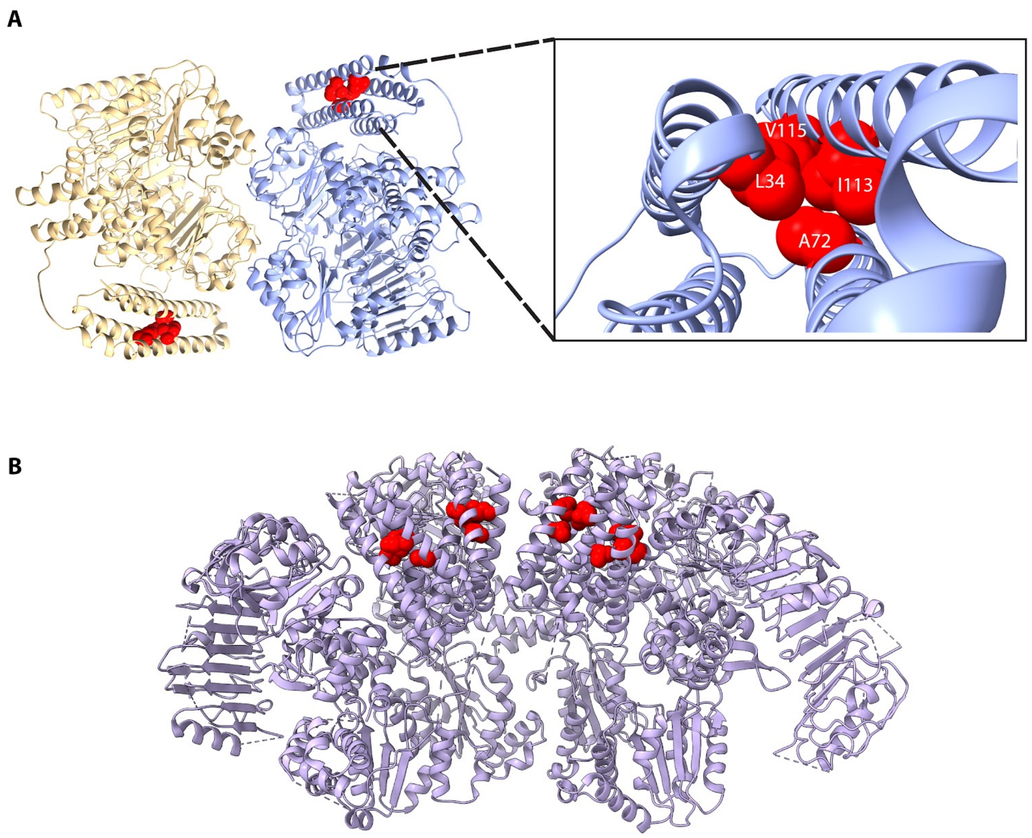


**Figure S4. The hydrophobic core of the CC domain in the NRC2 dimer and the resistosome**. (A) The hydrophobic core of the CC domain in the NRC2 homodimer (PDB: 8RFH), where the CC domain was predicted using AlphaFold3. Left panel: A side view of the α-helix barrel within the CC domain. (B) The position of the hydrophobic core in the hexameric NRC2 resistosome (PDB: 9FP6). Only four of the six NRC2 protomers are shown for clarity, with hydrophobic core residues highlighted in red.

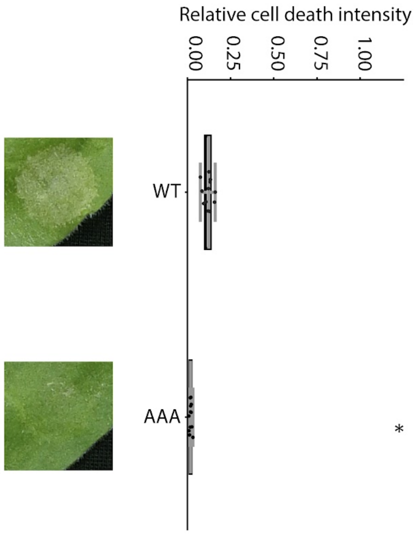


**Figure S5. NRC4$^{AAA}$ showed no cell death at 2 dpi compared to wild-type NRC4.** Cell death assay of wild-type and AAA variant of NRC4. The NRC4 and NRC4$^{AAA}$ were co-expressed with the Rpi-blb2 and AVRblb2 in *nrc2/3/4_KO N. benthamiana* leaves. The phenotype was recorded at 2 dpi. The cell death intensity was recorded by using UVP. The central line in each box plot represents the median cell death intensity, with box edges indicating the 25th and 75th percentiles. The whiskers extend to the most extreme data points no more than 1.5 x of the interquartile range. Statistical differences between wild-type and AAA were analyzed using the paired Wilcoxon signed rank test (*=p<0.01, **=p<0.005, ***=p<0.00001).

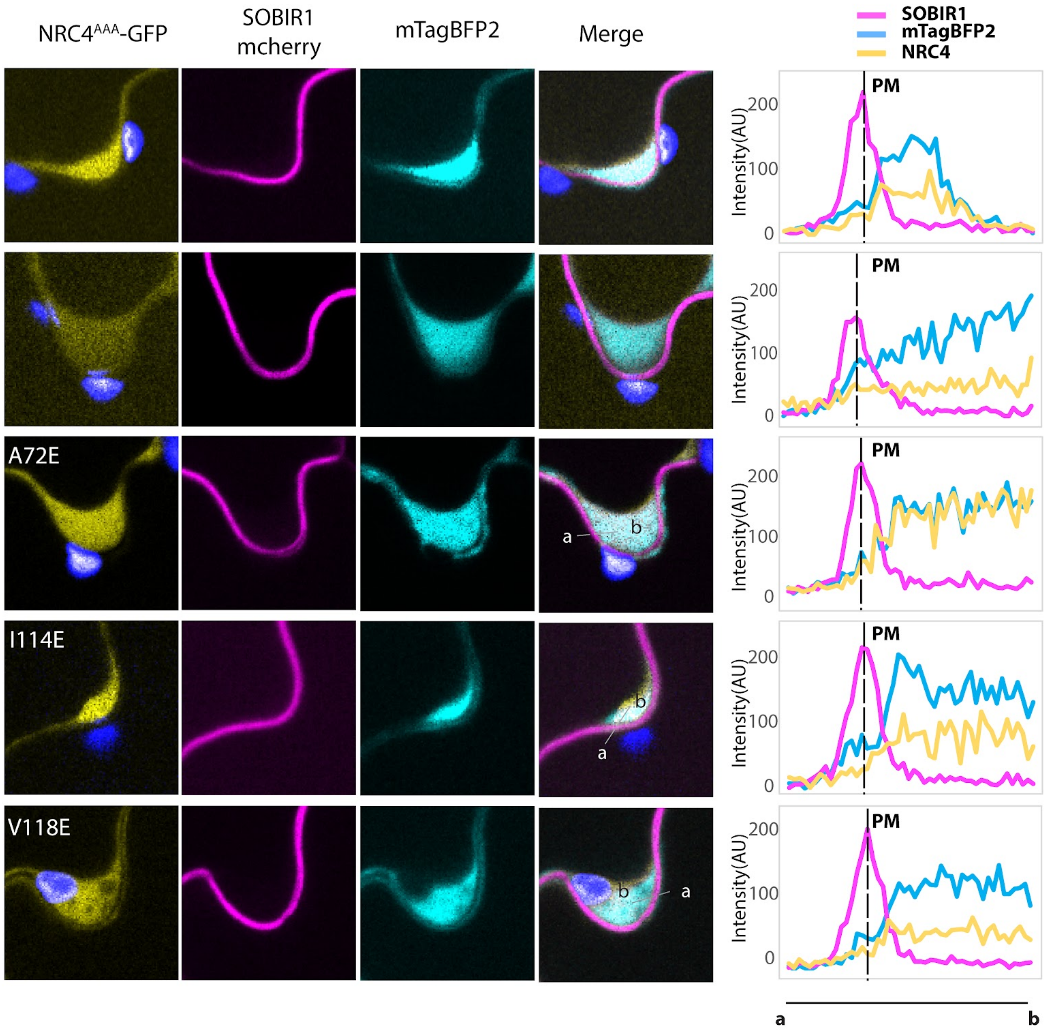


**Figure S6. Mutations on the hydrophobic core do not affect NRC4 subcellular localization in the resting state.** NRC4$^{AAA}$ and NRC4 hydrophobic core mutants were transiently co-expressed with Rpi-blb2, SOBIR1-mCherry (plasma membrane marker), and mTagBFP2 (cytosol marker) in *nrc2/3/4_KO N. benthamiana* leaves. Images were captured at 2 days post-infiltration (dpi) using confocal microscopy. The region exhibiting slight vacuole shrinkage was selected for fluorescence intensity measurements. The lines on the overlay panel indicate the area used to measure the intensity of the BFP (468 nm), GFP (525 nm), and RFP (615 nm) channels. The blue signal corresponds to the chloroplast. Fluorescence intensity was measured from position “a” to “b” in each group, with arbitrary units (AU) representing the values detected by the confocal microscope.

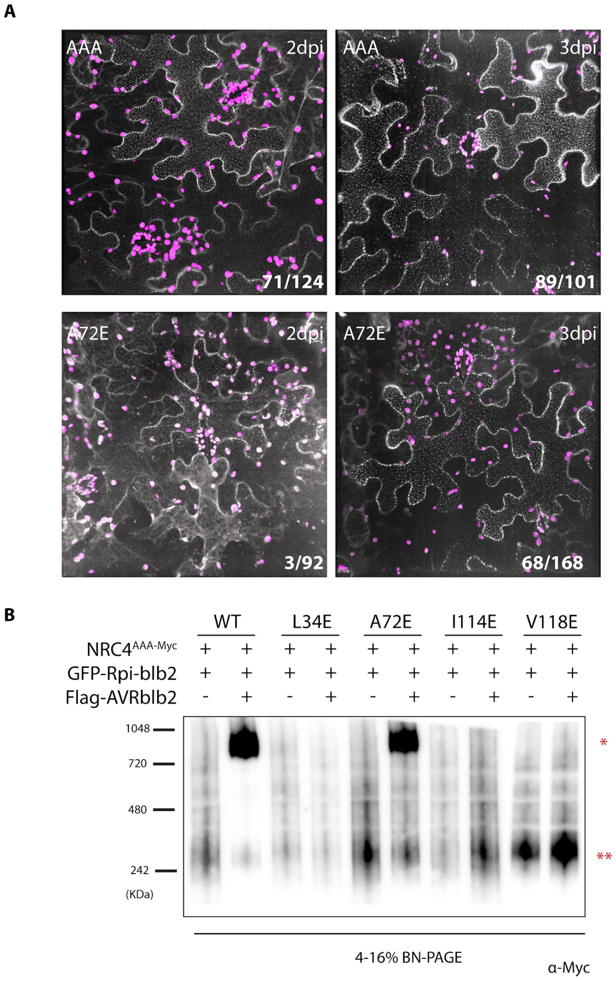


**Figure S7. The mutation on A72 delays the formation of the resistosome and puncta on the plasma membrane.** (A) Punctate formation of NRC4$^{AAA}$ and NRC4$^{AAA/A72E}$ at 2 and 3 dpi. NRC4$^{AAA}$ and NRC4$^{AAA/A72E}$, each fused to the C-terminal GFP, were transiently expressed along with Rpi-blb2 and AVRblb2 in *nrc2/3/4_KO N. benthamiana* leaves. Leaf discs from the infiltration areas were collected at 2 and 3 dpi. Confocal images were acquired as Z-stack projections of 40–50 slices. The day post-infiltration for each sample is shown in the top-right corner of each panel, and the number in the bottom-right corner indicates the proportion of cells exhibiting punctate formation out of the total cell count. The pink fluorescence corresponds to chloroplast autofluorescence. (B) BN-PAGE analysis of resistosome formation in NRC4$^{AAA}$ and NRC4$^{AAA/A72E}$. NRC4$^{AAA}$ and NRC4$^{AAA/A72E}$, tagged with a C-terminal 4×Myc epitope, were transiently co-expressed with Rpi-blb2 and AVRblb2 in *nrc2/3/4_KO N. benthamiana* leaves. Leaf discs were collected at 3 dpi and homogenized according to the protocol described in Materials and Methods. Total protein extracts were analyzed by BN-PAGE, and NRC4 variants were detected using an anti-Myc antibody. A single asterisk (*) denotes the resistosome, while a double asterisk (**) indicates the dimer.

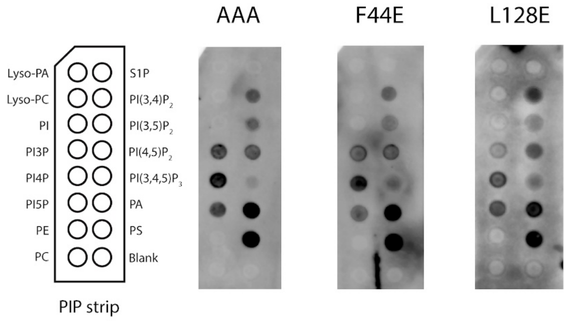


**Figure S8. Mutations of hydrophobic residues outside the hydrophobic core did not affect the association of the NRC4 CC domain with phospholipids.** Lipid binding assay for the CC domain of $NRC4^{AAA}$, $NRC4^{AAA/F44E}$, and $NRC4^{AAA/L128E}$. DNA sequences encoding the NRC4 CC domain (with a C-terminal Myc tag) were in vitro translated using the TNT Coupled Wheat Germ Extract System. The resulting protein extracts were applied to a PIP strip, and the NRC4 CC domain was detected using an anti-Myc antibody.

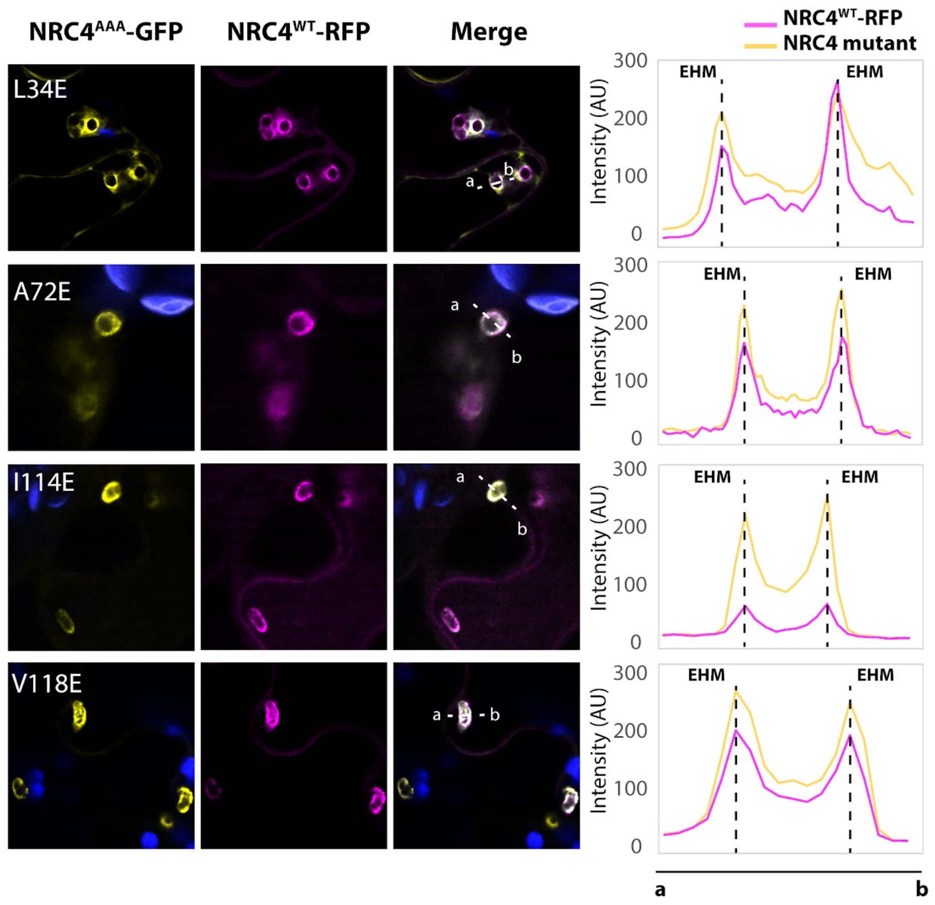


**Figure S9. Mutations of the hydrophobic core do not affect the NRC4 focal accumulation on the EHM.** The hydrophobic core mutants of NRC4 were tagged with GFP and transiently expressed with the wild-type NRC4-mCherry in the *nrc2/3/4_KO. N. benthamiana* leaf. After 1dpi, the zoospores of *P. infestans* were inoculated, and the leaves were imaged under confocal after 2 days post-inoculation. The lines marked on the overlay panel indicated the selected region for measuring the intensity of each channel. Fluorescence intensity was measured from position "a" to "b" in each group, and the values are presented as arbitrary units (AU).

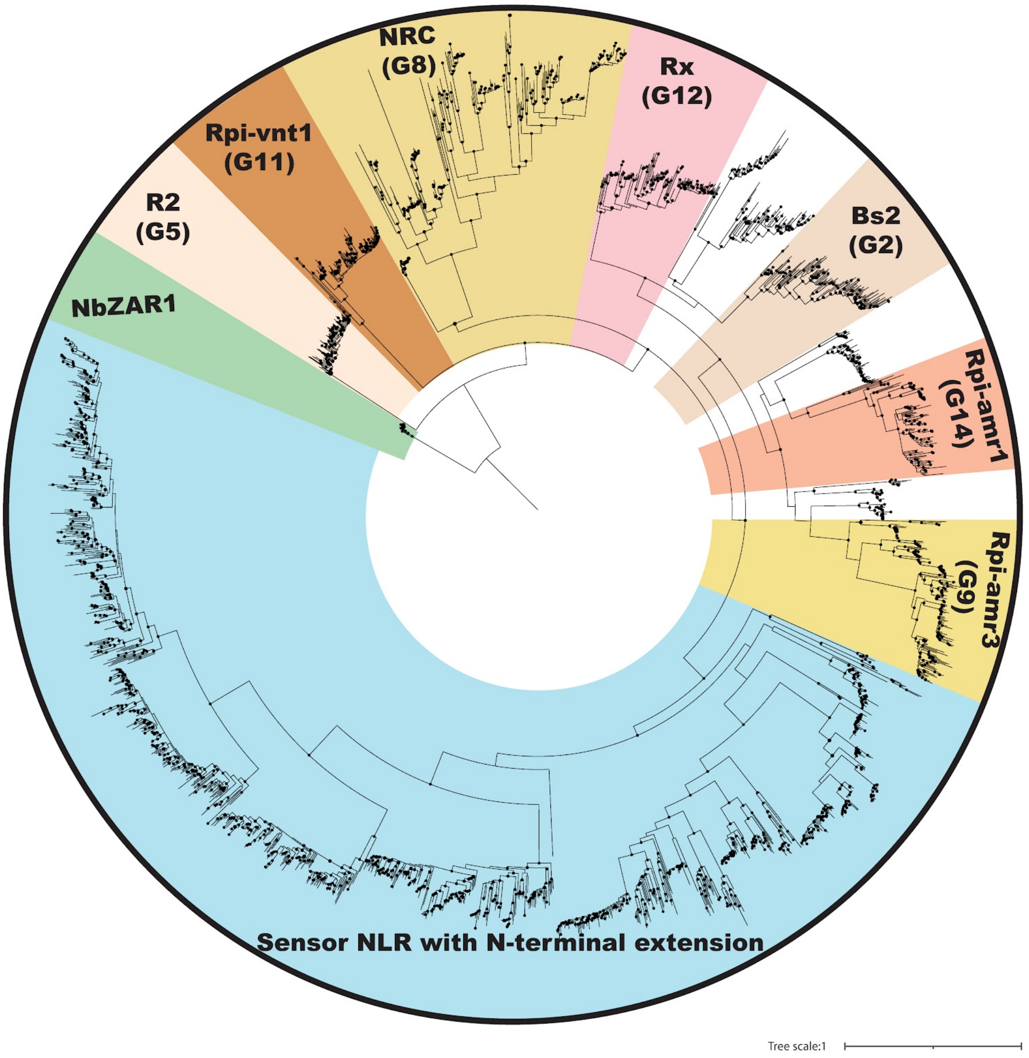


**Figure S10. Phylogenetic tree of the NRC superclade and selected singletons NLR groups (R2, Rpi-vnt1, and NbZAR1).** The detailed phylogenetic tree of Figure 6A, with each color denoting a distinct clade. A total of 2,797 NLR sequences from 22 representative solanaceous species are included. Black dots indicate nodes supported by bootstrap values above 70. The scale bar shows the evolutionary distance in amino acid substitutions per site.

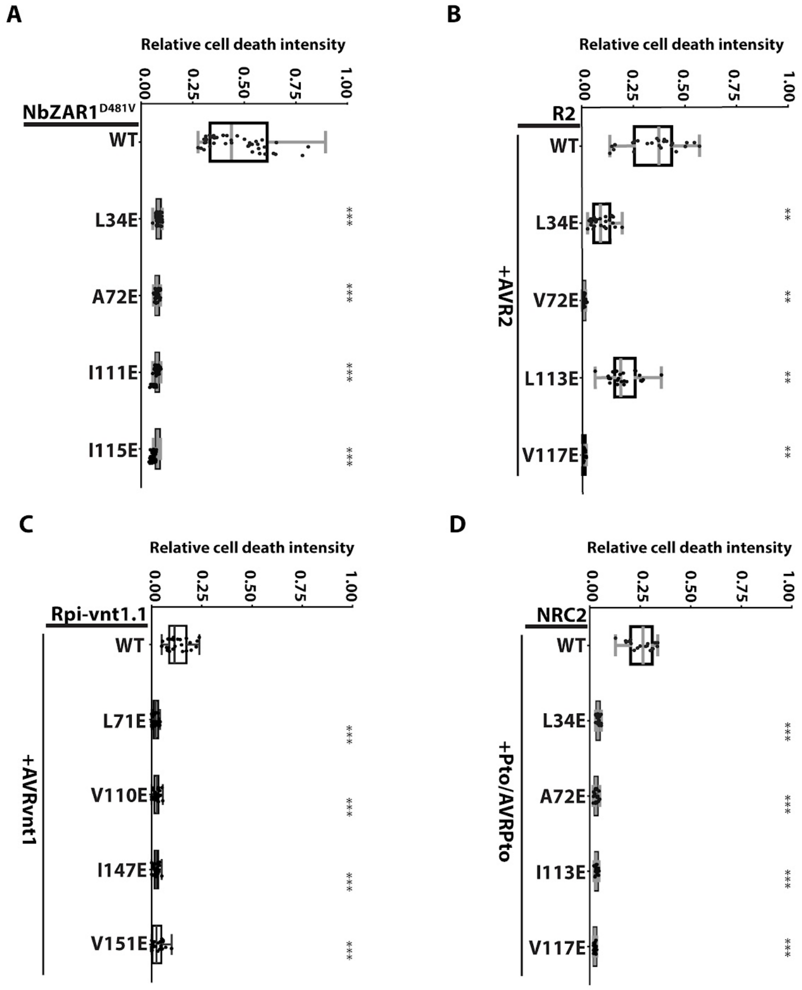


**Figure S11. Quantitative cell death result of NRC2, NRC3, R2, and NbZAR1 variants.** (A) NbZAR1 cell death assay. The autoactive variants of wild-type and hydrophobic core mutants of NbZAR1 were transiently expressed in wild-type *N. benthamiana* leaves. Cell death results were collected at 3 dpi. (B) Wild-type and hydrophobic core mutants of R2 were transiently expressed with the corresponding effector AVR2 in wild-type *N. benthamiana* leaves. Cell death results were collected at 3 dpi. (C) Wild-type and hydrophobic core mutants of Rpi-vnt1.1 were transiently expressed with the corresponding effector AVRvnt1 in wild-type *N. benthamiana* leaves. Cell death results were collected at 3 dpi. (D) NbNRC2 cell death assays. Wild-type or the hydrophobic core mutants NRC2 were transiently expressed with the serine/threonine kinase Pto and its corresponding effector AvrPto in *nrc2/3/4_KO N. benthamiana* leaves. Cell death intensity and phenotypes were recorded at 6 dpi. Cell death intensity was recorded using the UVP ChemStudio system. In each box plot, the central line represents the median cell death intensity, while the box edges indicate the 25th and 75th percentiles. Whiskers extend to the most extreme data points within 1.5 times the interquartile range. Statistical differences between wild-type and variant samples were analzyed using the paired Wilcoxon signed-rank test (*p < 0.01, **p < 0.005, ***p < 0.00001).

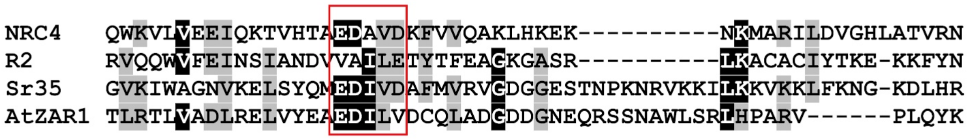


**Figure S12. The EDVID motif is absent in R2.** Sequence alignment of NbNRC4 R2, Sr35, and AtZAR1. The red rectangle indicates the EDVID motif in the amino acid sequence. Amino acids highlighted with a black background are identical in at least three reference sequences. In comparison, those with a gray background are similar but not identical in at least three reference sequences.

**Table S1. List of constructs used in cell death assays**

| Vector backbone | Promoter | protein name | Tag | $OD_{600}$ | Reference |
|---|---|---|---|---|---|
| pICH86988 | 35s | Sync-NRC4 wild-type | C-terminal Myc | 0.2 | (Wu *et al*, 2017) |
| pICH86988 | 35s | NRC4 variants | C-terminal Myc | 0.2 | This study |
| pK7WGF2 | 35s | Rpi-blb2 | N-terminal GFP | 0.2 | (Bozkurt *et al*, 2011) |
| pGWB12 | 35s | AVRblb2 | N terminal Flag | 0.1 | (Oh *et al*, 2009; Bozkurt *et al*, 2011) |
| pICH86988 | 35s | NRC2 Variants | C-terminal Myc | 0.2 | This study |
| pICH86988 | 35s | NRC3 Variants | C-terminal Myc | 0.2 | This study |
| pTFS40 | 35s | Pto | C-terminal HA | 0.2 | (Rathjen *et al*, 1999; de Vries *et al*, 2006) |
| pT50 | 35s | AvrPto | C-terminal Flag | 0.1 | (Rathjen *et al*, 1999; de Vries *et al*, 2006) |
| pICH86988 | 35s | NbZAR1 variants | C-terminal Myc | 0.2 | This study |
| pICH86988 | 35s | R2 variants | none | 0.2 | This study |
| pICH47751 | 35s | AVR2 | none | 0.1 | (Gilroy *et al*, 2011) |
| pICH86988 | 35s | Rpi-vnt1.1 variants | none | 0.2 | This study |
| pK7WG2 | 35s | AVRvnt1 | none | 0.1 | (Foster *et al*, 2009) |

**Table S2. List of constructs used in lipid binding assays**

| Vector backbone | Promoter | protein name | Tag |
|---|---|---|---|
| pICSL86900OD | T7 promoter | mCherry | C terminal Myc |
| pICSL86900OD | T7 promoter | $NRC4^{AAA}$ CC | C terminal Myc |
| pICSL86900OD | T7 promoter | $NRC4^{AAA/L34E}$ CC | C terminal Myc |
| pICSL86900OD | T7 promoter | $NRC4^{AAA/F44E}$ CC | C terminal Myc |
| pICSL86900OD | T7 promoter | $NRC4^{AAA/A72E}$ CC | C terminal Myc |
| pICSL86900OD | T7 promoter | $NRC4^{AAA/I114E}$ CC | C terminal Myc |
| pICSL86900OD | T7 promoter | $NRC4^{AAA/V118E}$ CC | C terminal Myc |
| pICSL86900OD | T7 promoter | $NRC4^{AAA/L128E}$ CC | C terminal Myc |

**Table S3. List of constructs used in cell biology assays**

| Vector backbone | Promoter | protein name | Tag | $OD_{600}$ | Reference |
|---|---|---|---|---|---|
| pICH86988 | 35s | $NRC4^{AAA}$ | C-terminal GFP | 0.2 | This study |
| pICH86988 | 35s | $NRC4^{AAA/L34E}$ | C-terminal GFP | 0.2 | This study |
| pICH86988 | 35s | $NRC4^{AAA/A72E}$ | C-terminal GFP | 0.2 | This study |
| pICH86988 | 35s | $NRC4^{AAA/I114E}$ | C-terminal GFP | 0.2 | This study |
| pICH86988 | 35s | $NRC4^{AAA/V118E}$ | C-terminal GFP | 0.2 | This study |
| pICH47751 | 35s | Rpi-blb2 | N-terminal Hellfire | 0.2 | (Huang *et al*, 2024) |
| pGWB12 | 35s | AVRblb2 | N-terminal Flag | 0.1 | (Tameling & Baulcombe, 2007) |
| pK7WG2 | 35s | SlSOBIR1 | C-terminal mCherry | 0.2 | (Li *et al*, 2021) |
| pICH47792 | 35s | mTagBFP2 | Myc | 0.2 | This study |

**Table S4. List of constructs used in BN-PAGE assays**

| Vector backbone | Promoter | protein name | Tag | $OD_{600}$ | Reference |
|---|---|---|---|---|---|
| pICH86988 | 35s | $NRC4^{AAA}$ | C-terminal Myc | 0.2 | This study |
| pICH86988 | 35s | $NRC4^{AAA/L34E}$ | C-terminal Myc | 0.2 | This study |
| pICH86988 | 35s | $NRC4^{AAA/A72E}$ | C-terminal Myc | 0.2 | This study |
| pICH86988 | 35s | $NRC4^{AAA/I114E}$ | C-terminal Myc | 0.2 | This study |
| pICH86988 | 35s | $NRC4^{AAA/V118E}$ | C-terminal Myc | 0.2 | This study |
| pK7WGF2 | 35s | Rpi-blb2 | N-terminal GFP | 0.2 | (Bozkurt *et al*, 2011) |
| pGWB12 | 35s | AVRblb2 | N-terminal Flag | 0.1 | (Bozkurt *et al*, 2011) |

**Table S5. List of constructs used in cell fractionation assays**

| Vector backbone | Promoter | protein name | Tag | $OD_{600}$ | Reference |
|---|---|---|---|---|---|
| pICH86988 | 35s | $NRC4^{AAA}$ | C-terminal Myc | 0.2 | This study |
| pICH86988 | 35s | $NRC4^{AAA/L34E}$ | C-terminal Myc | 0.2 | This study |
| pICH86988 | 35s | $NRC4^{AAA/A72E}$ | C-terminal Myc | 0.2 | This study |
| pICH86988 | 35s | $NRC4^{AAA/I114E}$ | C-terminal Myc | 0.2 | This study |
| pICH86988 | 35s | $NRC4^{AAA/V118E}$ | C-terminal Myc | 0.2 | This study |
| pK7WGF2 | 35s | Rpi-blb2 | N-terminal GFP | 0.2 | (Bozkurt *et al*, 2011) |
| pGWB12 | 35s | AVRblb2 | N-terminal Flag | 0.1 | (Bozkurt *et al*, 2011) |
| pICH47792 | 35s | mTagBFP2 | C-terminal Myc | 0.2 | This study |

**Table S6. List of constructs used in focal accumulation assays**

| Vector backbone | Promoter | protein name | Tag | $OD_{600}$ | Reference |
|---|---|---|---|---|---|
| pH7GWR2 | 35s | NRC4 wild-type | C-terminal GFP | 0.2 | (Duggan *et al*, 2021) |
| pICH86988 | 35s | $NRC4^{AAA}$ | C-terminal GFP | 0.2 | This study |
| pICH86988 | 35s | $NRC4^{AAA/L34E}$ | C-terminal GFP | 0.2 | This study |
| pICH86988 | 35s | $NRC4^{AAA/A72E}$ | C-terminal GFP | 0.2 | This study |
| pICH86988 | 35s | $NRC4^{AAA/I114E}$ | C-terminal GFP | 0.2 | This study |
| pICH86988 | 35s | $NRC4^{AAA/V118E}$ | C-terminal GFP | 0.2 | This study |
| pGWB555 | 35s | mRFP | non | 0.2 | (NAKAGAWA *et al*, 2007) |
